# PRO-LDM: Protein Sequence Generation with a Conditional Latent Diffusion Model

**DOI:** 10.1101/2023.08.22.554145

**Authors:** Sitao Zhang, Zixuan Jiang, Rundong Huang, Shaoxun Mo, Letao Zhu, Peiheng Li, Ziyi Zhang, Emily Pan, Xi Chen, Yunfei Long, Qi Liang, Jin Tang, Renjing Xu, Rui Qing

## Abstract

Deep learning-driven protein design holds enormous potential despite the complexities in sequences and structures. Recent developments in diffusion models yielded success in structure design, but awaits progress in sequence design and are computationally demanding. Here we present PRO-LDM: an efficient framework combining design fidelity and computational efficiency, utilizing the diffusion model in latent space to design proteins with property tuning. The model employs a joint autoencoder to capture latent variable distributions and generate meaningful embeddings from sequences. PRO-LDM (1) learns representations from biological features in natural proteins at both amino-acid and sequence level; (2) generates native-like new sequences with enhanced diversity; and (3) conditionally designs new proteins with tailored properties or functions. The out-of-distribution design enables sampling notably different sequences by adjusting classifier guidance strength. Our model presents a feasible pathway and an integratable tool to extract physicochemical and evolutionary information embedded within primary sequences, for protein design and optimization.

## Introduction

Proteins are miniscule molecular machines that undertook a vast range of indispensable biological functions and sustain the life of organisms. Yet native proteins only occupy a limited proportion of the vast sequence space. Protein design is the process where researchers modify native proteins or construct sequences from scratch to achieve tailored properties or functions by accurately placing amino acids (*1*), to obtain novel biomolecules more suitable for specific applications. Protein design has propelled breakthroughs in pharmaceutical (*2–4*), material (*5, 6*), and environmental conservation fields (*7*). Common protein design strategies include rational design (*8*), site-directed mutagenesis, directed-evolution (*9–11*), semi-rational design (*12–14*), and computational assisted design (*15–17*). Rational design pioneered the exploration of proteins’ sequence-structure-function relationships through manually altering amino acids at specific sites in a hypothesis-driven strategy, often achieved through site-directed mutagenesis (*18*). Yet the approach is often limited by the availability of prior knowledge on the specific target of modification. Directed-evolution resembles nature’s evolutional process and involves multiple rounds of amino acid mutation, screening, and selection. It iterates in a semi-guided manner to approach target protein performances (*18*), but is typically labor intensive and still evolves around the sequence spaces adjacent to native proteins. On the other hand, computational design methods leverage the ever-expanding protein databases to facilitate more precise design of sequences and structures, while drastically reducing the reliance on high-throughput experimental screenings (*19*). The advent of deep learning-based algorithms further expands the prospect not only in protein design but across the entire realm of biology.

The most recent development that changed the paradigm in molecular biology research is the AlphaFold2 algorithm, which combines attention-based neural networks with evolutional conservation information in proteins to predict their 3D structures from sequences with exceptional accuracy (*20*). It significantly outperformed traditional computational prediction methods (homology modeling, *de novo* modeling, *etc.*), and greatly supplemented the availability of protein crystallography database while reducing the costs and time. The advent of AlphaFold2 and other deep learning-based structure prediction algorithms such as RoseTTAFold represents a pivotal milestone in the application of artificial intelligence in molecular biology.

Since structure prediction and protein design are two sides of the same coin, researchers quickly started to explore the use of these algorithms in designing novel protein species. Deep generative models have garnered significant attention for this purpose. Their excellent track-records in language and image processing showed capability of learning from vast unlabeled datasets, generating meaningful data representations, and producing new samples that resemble the distributions of training data, all of which are transplantable to protein design tasks. Groundbreaking works like DALL-E 2 (*21*) and GPT (*22–24*) highlight the enormous potential in generating diverse and realistic data. As the latest state-of-the-art (SOTA) generative models, diffusion models (*25–28*) have emerged as a widely applied and highly effective approach in multiple domains (*29–34*) with ability to sample complex distributions, that have integrative and controllable refinement processes and generate high-fidelity and robust species in diverse sequence and structural spaces. Lee *et al.* has reported ProteinSGM, a score-based generative model capable of designing new proteins with conformational folds not presenting in the training set (*35*). A conditional diffusion model was then built and tested on three design cases to generate structures to be inserted into masked sequences that corresponded well to native conformations with a mean pLDDT (per-residue Local Distance Difference Test) of >80. Watson *et al*. developed RFdiffusion through fine-tuning the RoseTTAFold on protein structure denoising for protein main chain generation (*36*). This model was demonstrated in various applications, including unconditional protein design, binder design, oligomer design, enzymic and metal-binding site design, *etc*.

Whilst diffusion models have made notable progress in protein structure design, its potential in sequence design still has much to be explored (*37*). The application of diffusion models also involves multiple denoising iterations with high demand in computational power (*25*). Here we present PRO-LDM (Protein Sequence Generation with conditional Latent Diffusion Models), a multi-task learning algorithm that combines sequence generation and fitness prediction in a single framework. Our architecture adopts a diffusion module in the latent space for sequence construction, which may also be integrated with alternative encoders and pre-trained models for enhanced scalability and adaptivity. PRO-LDM is capable of extracting biological representations from both single amino acids and whole protein sequences, as demonstrated by latent space visualization. The latent variable distributions are captured to generate meaningful embeddings for unconditionally designing new sequences with native-like characteristics and high diversity, while global amino acid relationships are also reproduced. The conditional design results in new proteins within target fitness ranges, that might be used in protein property and functional optimizations. Moreover, PRO-LDM also shows advantages over single-task models in reduced computational time and more cost-effective maintenance. Our model presents a modular, combinatorial new tool to extract biological information within protein sequences and generate novel species with target features.

## Results

### 1. The architecture of PRO-LDM

PRO-LDM was based on a jointly trained autoencoder (JT-AE) with a latent conditional diffusion (LCD) module to learn fundamental patterns embedded within protein sequences. The model can be tailored to conduct both unconditional and conditional protein sequence design with enhanced diversity. A fitness label was assigned to define the respective property or function in a given set of proteins. PRO-LDM can also conditionally generate protein sequences towards a target label and predict their fitness accordingly, when trained with sequence dataset with label values. If unlabeled datasets were used or the labels were uniformly set to 0, PRO-LDM conducted unsupervised learning to unconditionally generate protein sequence sets with similar properties to the training sets.

#### JT-AE

JT-AE was the fundamental structure of ReLSO (*38*) and a combination of supervised and unsupervised learning in an autoencoder framework. It consisted of a transformer-based encoder, a convolutional neural network (CNN)-based decoder and a multilayer perceptron (MLP)-based regressor in parallel. The output of the encoder was subjected to dimension reduction through a bottleneck module composed of fully connected layers, resulting in latent variable *z*. The collection of all *z* values constituted the latent space. Latent variable *z*, along with the labels representing sequence fitness, were simultaneously passed to the diffusion module to simulate the distribution in the latent space, as elaborated below.

#### Latent conditional diffusion model

LDM learned the data distribution in the latent space from sequences and captured their characteristics with different fitness. It employed a UNet (*39*) as the neural network backbone and utilized ancestral sampling for the generation process like denoising diffusion probabilistic models (DDPM) (*25*).

During the training phase, given an input sequence *x*, the encoder *f*_*θ*_ encoded *x* into a latent representation *z* = *f*_*θ*_ (*x*), and the decoder *g*_*θ*_ reconstructed the sequence from the latent variable, giving *x̃* = *g*_*θ*_ (*z*) = *g*_*θ*_ (*f*_*θ*_ (*x*)). In the latent space, we divided the diffusion process into *T* steps and added Gaussian noise according to a variance schedule *β*_1_, … , *β*_*T*_:

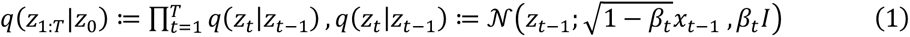

where *z*_0_∼*p*(*z*_0_). The reverse process could be defined as a Markov chain, starting at *p*_*θ*_ (*z*_*T*_) = *N*(0, *I*):

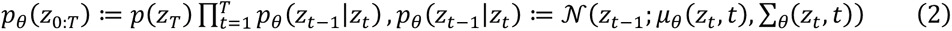

*β*_*t*_ was kept constant as a hyperparameter, where 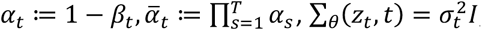, and 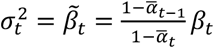. Consequently, we could sample 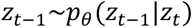 using the following equation:

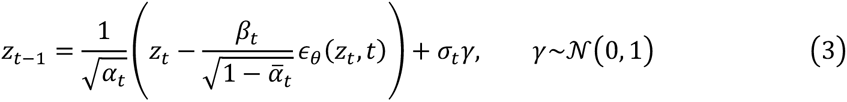

where *∈*_*θ*_ was a function approximator designed to predict *∈* from *z*_*t*_. In the case of conditional sampling, the latent variable *z* was drawn along with class label *c*, so that the function approximator was changed into *∈*_*θ*_ (*z*_*t*_ , *t*, *c*). We jointly trained an unconditional diffusion model *p*_*θ*_ (*z*) parameterized through a function approximator *∈*_*θ*_ (*z*_*t*_ , *t*), along with the conditional model *p*_*θ*_ (*z*|*c*) parameterized through *∈*_*θ*_ (*z*_*t*_ , *t*, *c*) , by randomly setting *c* to the unconditional class identifier ⌽ with a probability *p_uncond_* as a hyperparameter:

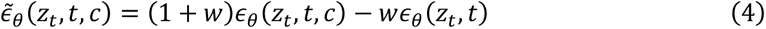

where *w* was a hyperparameter that controlled the strength of the classifier-free guidance. Consequently, we could train the latent diffusion model using the following equation:

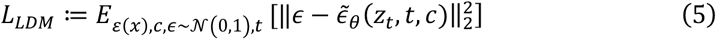

Although capable of generating high-fidelity samples with diverse data distributions, diffusion models were computationally expensive. To overcome this issue, our model utilized the diffusion process in the latent space to reduce the dimensionality of data, compared to other models that used diffusion all through the whole process. This architecture can accelerate the sequence generation speed and improve model efficiency. Moreover, we circumvented the need for classifier-guided diffusion, which required an additional pretrained classifier that may further increase model complexity. Instead, our classifier-free guided diffusion combined conditional and unconditional diffusions for joint training, striking a good balance between model complexity and computational costs, without harming the quality in sequence generation.

The model was trained by minimizing the loss below.

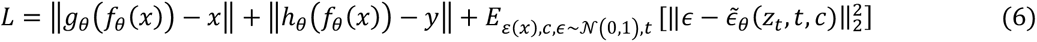

The first and second term gauged the loss of JT-AE, where *f*_*θ*_ represented the encoder, *g*_*θ*_ represented the decoder, *h*_*θ*_ represents the regressor, *x* was the input sequence and *y* was the corresponding fitness. The third term measured the loss of the latent diffusion model which was described above in detail.

### 2. PRO-LDM learns representations of protein sequences

Our model was first trained unconditionally on protein datasets without associated fitness values, to capture inherent property or function representations in sequences. PRO-LDM was able to generate new variants of proteins by learning information embedded solely in original sequences. The model was trained respectively on three datasets, including two datasets on the homologs of bacterial luciferase procured from InterPro (IPR011251), and one dataset on a family of bacterial MDH enzymes (EC1.1.1.37). The luciferase protein sets were divided into two different forms, namely, Luciferase_MSA and Luciferase_RAW, to determine how multiple sequence alignment (MSA) affect the learning efficacy and generative performance of the model. MSA was performed for all data in Luciferase_MSA, where a gap character was assigned to all unfilled positions during training. Sequences in Luciferase_RAW were aligned to the N-terminus, while additional padding characters were added to the C-terminal end to ensure all sequences were of the same length.

The propensity and distribution of 20 amino acids in a given sequence defined the structure and function for proteins. Hence, it was essential for a learning algorithm to capture the intricate biochemical attributes of amino acids. In general, amino acids with similar side chain structures and physicochemical properties were more likely to strongly correlated during the learning process that surpassed those among amino acid not alike. Therefore, we first extracted embeddings of amino acids from single sequences and calculated their correlations. An example of this was shown in fig. 2A and table S1 where stronger correlation between individual amino acids with similar properties were revealed. The embeddings of corresponding amino acids were further integrated from the entire dataset and processed through the principal component analysis (PCA) dimension reduction algorithm to visualize in the 2D space. Fig. 2B showed results for all three datasets, where amino acids more alike were positioned near each other, such as charged acidic and basic amino acids. In contrast, amino acids with distinct biochemical properties were spaced further apart, such as nonpolar and polar amino acids. The results demonstrated that our model was able to learn the characteristics of amino acids solely from their appearances in sequences.

**Fig. 1.**
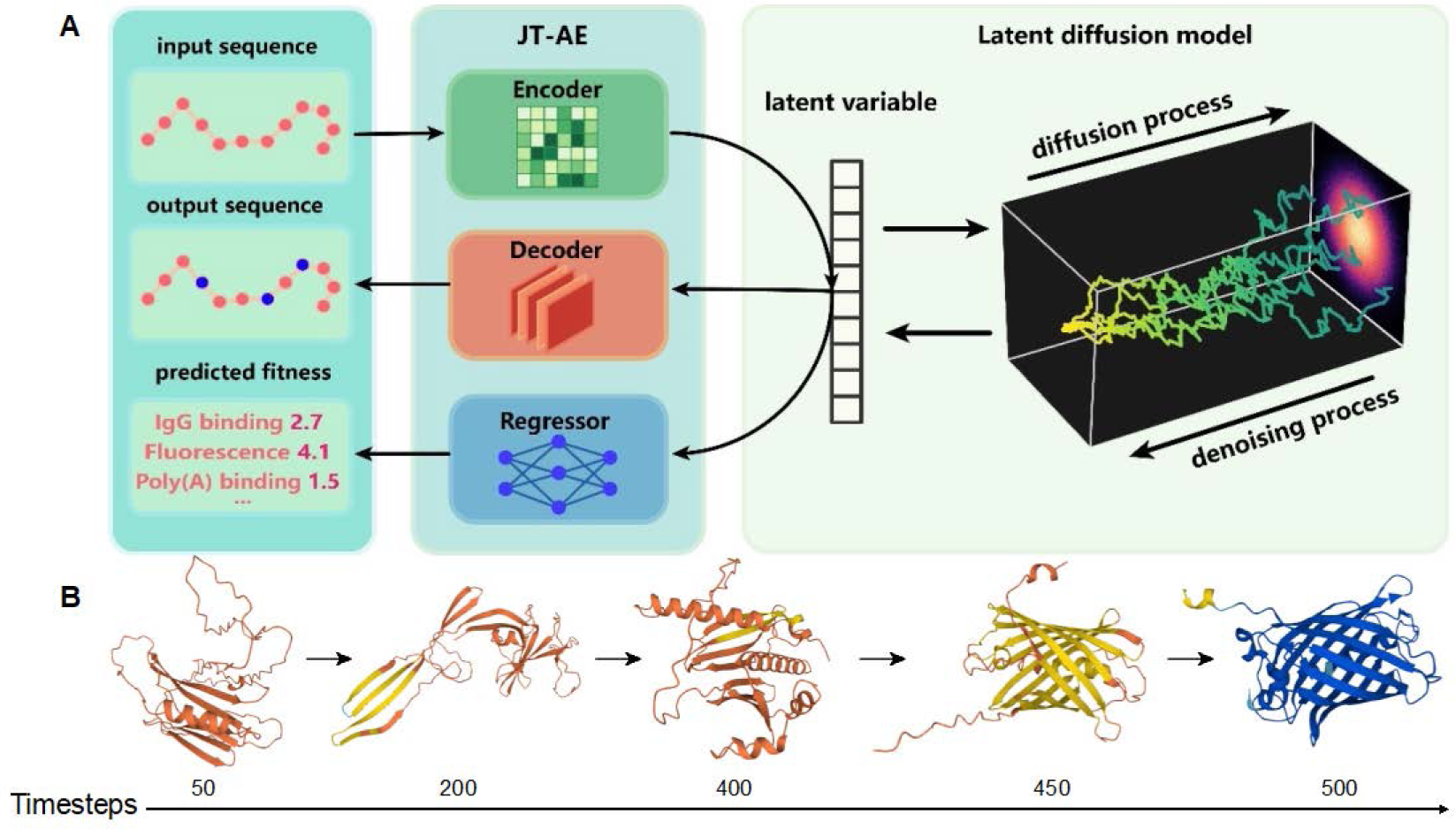
Overview of the PRO-LDM. **(A)** the PRO-LDM architecture. In the training stage, input sequences were mapped into the latent space via a transformer-based encoder. A latent diffusion model was then applied to capture the distribution of the latent space. Finally, the latent variables were used to reconstruct the sequence via a CNN-based decoder and simultaneously predict the fitness via an MLP-based regressor. In the sampling stage, the latent variables of new sequences were generated via the denoising process of LDM starting from a simple noise distribution. The output sequences and predicted fitness were obtained using the decoder and regressor, respectively. **(B)** 3D-structure iterations of a generated protein during the denoising process for GFP (green fluorescent protein) colored to the pLDDT value (deep blue: pLDDT > 90; light blue: 90 > pLDDT > 70; yellow: 70 > pLDDT > 50). We selected several time steps in the denoising process trained on the GFP dataset and decoded the latent variables into sequences. AlphaFold2 was implemented to predict the 3D structures to visualize the modeling process.

**Fig. 2.**
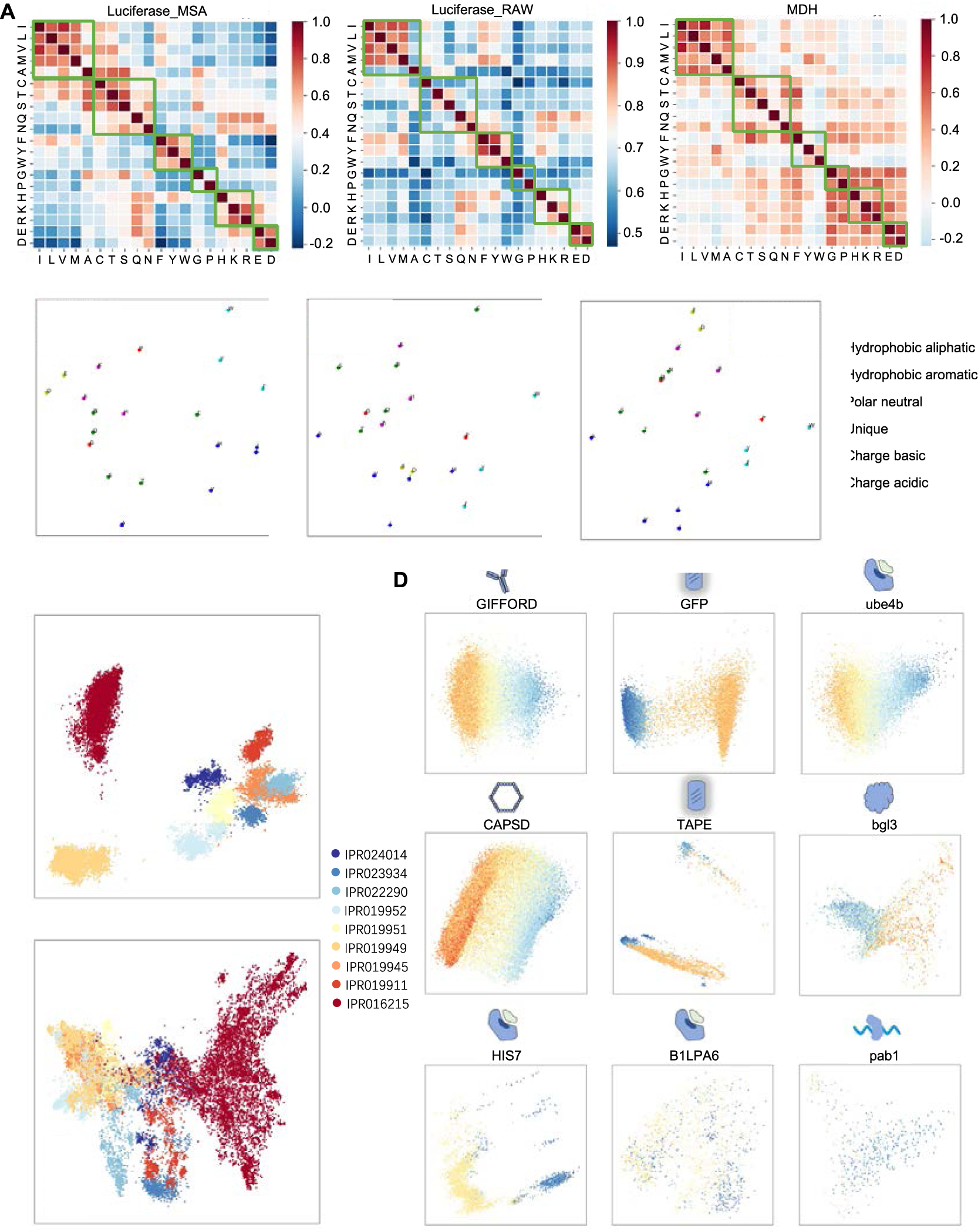
Protein representations at amino acid and sequence level. **(A)** Pearson’s correlation between amino acid types in randomly selected sequences. For each output sequence, a 2D matrix was generated to represent the likelihood of different amino acids occurring at each residue position. Pearson’s correlation was then calculated to measure the relationship between each amino acid pair. Biochemically related amino acids exhibited higher correlation. Amino acid pairs within the same green border shared similar characteristics. Dataset from left to right: Luciferase_MSA, Luciferase_RAW, MDH. **(B)** Average latent space representations of amino-acid characteristics learned by PRO-LDM. (n = 20 amino acids, dataset from left to right: Luciferase_MSA, Luciferase_RAW, MDH.) **(C)** Organization of latent space reflecting subfamilies of Luciferase. Visualizations illustrated the latent representation of sequences in Luciferase_MSA (top) and Luciferase_RAW (bottom), projected onto the first two principal components and colored by sub-family annotations derived from InterPro. Only sequences belonging to the 9 largest subfamilies were shown. **(D)** Latent space representation of labeled protein sequences. The latent embeddings of 9 labeled datasets learned by PRO-LDM were displayed. The protein sequence representations were visualized by PCA, and each point was colored according to its corresponding fitness value. From blue to orange: high fitness to low fitness.

Using the luciferase dataset as an example, we evaluated the efficiency of our model on learning comprehensive representations of protein sequences. Luciferase proteins were classified into different subfamilies based on their diverse folds, the information of which was extracted from InterPro and the 9 largest subfamilies were selected for our analysis. Sequence embedding and family information were visualized in fig. 2C, where results for both Luciferase_MSA and Luciferase_RAW were shown. Apparent clustering was observed for sequences belonging to the same subfamily in both training sets. The processing on MDH, Luciferase_MSA and Luciferase_RAW training sets clearly showed that our model not only captured embedded characteristics of amino acids solely from their positional appearances within sequences, but also grasped attributes of proteins as a whole related to their properties and functions, which laid the foundation for subsequent design tasks.

High-throughput molecular biology techniques such as deep mutational scanning (DMS), directed-evolution, saturation mutagenesis, and random mutagenesis were common tools to reveal the intricate relationship between genotype and phenotype (*40*). These techniques yielded large amounts of mutational data showing how genetic changes affected the function of proteins and living organisms. In our study, 9 mutation datasets were used to train PRO-LDM. These datasets included varieties of sequences with equal or unequal lengths, as well as both indels (insertions/deletions) and amino acid substitutions (Methods; Table S2). Dimension reduction of sequence embeddings was performed by PCA into 2D space, where fitness values were presented by different colors. As shown in fig. 2D, latent space visualizations of most datasets exhibited global organization to fitness, which laid the foundation for conditional sequence generation of functional variants with tailored properties. PRO-LDM was also capable of predicting the fitness for protein sequences. The results were comparable to JT-AE in 9 datasets (table S3).

### 3. PRO-LDM learns intrinsic characteristics in sequences and designs variants similar to natural proteins

The core objective of a generative model was to create new data that followed the same distribution as original data. From the aspect of protein design, this expanded the known sequence space, and generated a greater variety of proteins with properties or functions that resembled or surpassed native species. PRO-LDM was designed to perform this task based on the training datasets when a fitness label was not assigned. Due to notable accomplishments from VAE-based models in capturing embedded information that distinguished protein sequences and generating native-like sequences in the target latent space, it was selected for comparison with PRO-LDM (*41*).

The progress of training was monitored by comparing the identity between generated and natural sequences through calculating the proportion of identical residues in both sets. Sixty-four sequences were generated in every 50 epochs. The identity between generated and natural sequences were observed to increase simultaneously with the training steps (fig.3A, top). For the MDH dataset, PRO-LDM resulted in higher median identity compared to VAE at the same epochs and achieved a higher final range after convergence (fig.3A, left top *vs.* down). For the luciferase_MSA dataset, the identity of PRO-LDM was lower than VAE in the first 50 epoch, but reached a higher level after convergence (fig.3A, medium top *vs.* down). For the luciferase_RAW dataset, the performance of VAE model was relatively poor, with a consistently low level of identify (the highest value being less than 40%). In contrast, PRO-LDM achieved a final identity of 90% or higher, showing significantly enhanced learning capability (fig.3A, right top *vs.* down). The comparison between the two luciferase datasets also suggested that training PRO-LDM with data subjected to MSA led to faster convergence and helped to generate sequences more similar to natural proteins. Therefore, integrating evolutionary information in the training data could further enhance the learning efficiency of PRO-LDM.

**Fig. 3.**
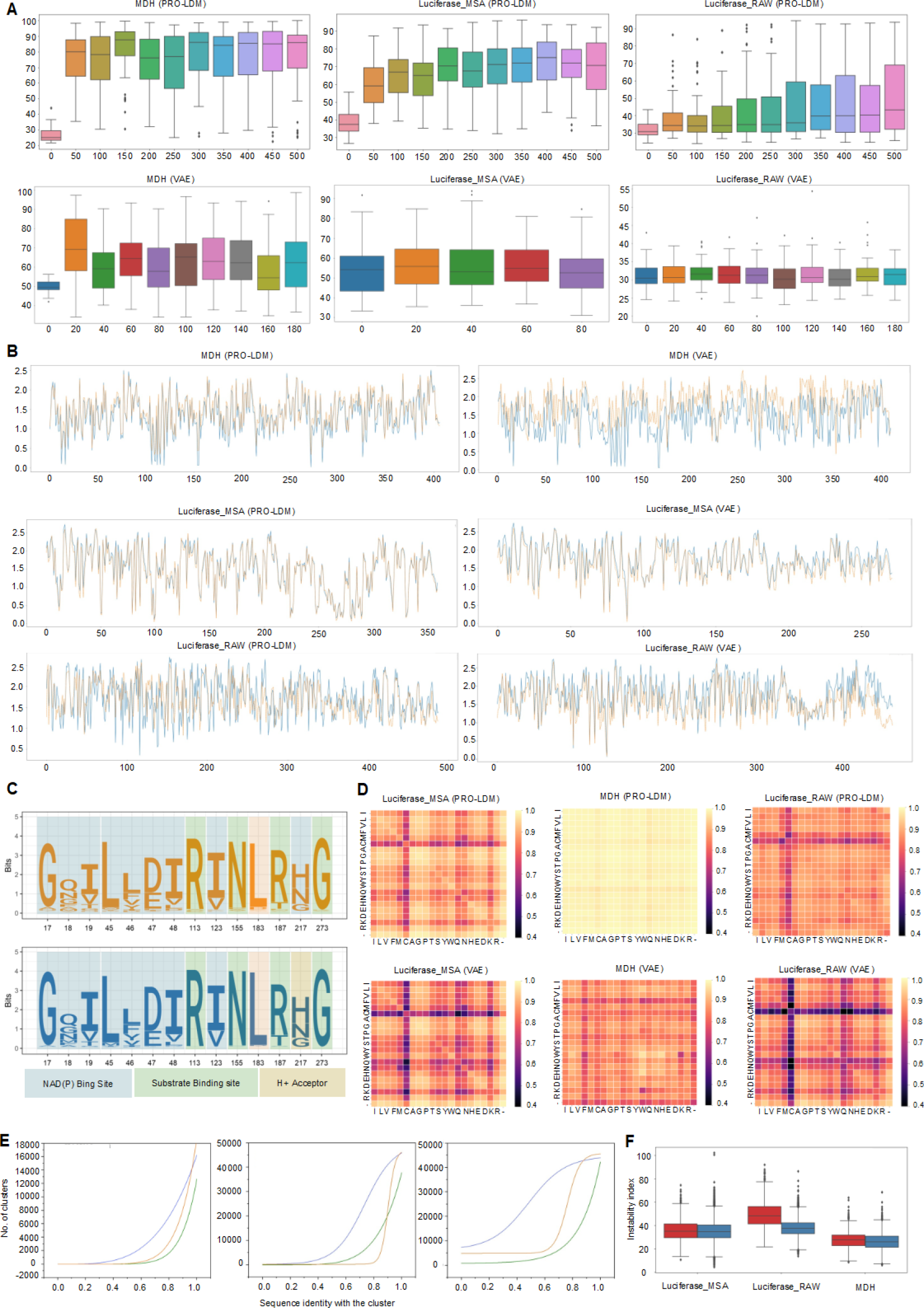
Unconditional protein design by PRO-LDM. **(A)** Sequence identity of 64 generated sequences to the nearest natural sequence from training data at different training iterations. X axis: training epoch; Y axis: sequence identity between generated sequence and best match sequence in training set. **(B)** Positional variability evaluation of PRO-LDM or VAE generated sequences (orange) versus natural sequences (blue). X axis: amino acid position; Y axis: Shannon entropy. **(C)** A sequence logo figure of key conserved positions in the MSA of MDH datasets. **(D)** Amino acid pairwise correlations of generated and natural sequences. Each point on the map represented the correlation of amino-acid pair frequencies between the natural sequences of Luciferase_MSA, Luciferase_RAW, MDH and those generated by PRO-LDM or VAEs. A high correlation denoted that the same pairwise long-distance amino-acid interactions were found as in natural sequences. **(E)** Comparison of sequence diversity for natural and generated sequences in three datasets (left: MDH; mid: Luciferase_MSA; right: Luciferase_RAW). **(F)** Comparison of sequence stability for natural and generated sequences in three datasets (left: Luciferase_MSA; mid: Luciferase_RAW; right: MDH).

Amino acid conservations in proteins were associated with critical structural and functional motifs due to nature’s selection process (*42, 43*). The tendency of amino acids staying unchanged at specific positions often represented key evolutional information passed on over a long period of time, the disruption of which might cause significant conformational changes in proteins that disabled them from carrying out original functions. Such positional variability in sequences can be determined by calculating Shannon entropy, for each site in the MSAs of generated and training sets (*44, 45*). Sequences generated by PRO-LDM exhibited highly similar Shannon entropy profiles compared to those in training sets (fig. 3B, table S4), indicating that critical residue positions along with evolutional conservation patterns were learnt from the natural sequences by PRO-LDM. In MDH dataset, our algorithm demonstrated remarkably superior performance over the VAE model, which reduced the positional entropy error (m.s.e) of VAE-generated sequences by over 11-fold (fig. 3B top). In the two luciferase datasets, the positional variabilities observed in PRO-LDM generated sequences and natural sequences exhibited a high degree of similarity (with an overall correlation coefficient exceeding 0.76), and were slightly superior to that of the VAE model (fig. 3B medium and bottom). Additionally, the high similarity of Shannon entropy between generated sequences and natural proteins in MDH and luciferase_RAW indicated the capture of intrinsic evolutionary patterns even without alignment information.

Enzymes in the MDH dataset need to bind both its substrate and the NAD^+^ cofactor to carry out catalytic functions. Hence, we analyzed the functionality of generated sequences by predicting functional sites via InterPro and marking them in the logo figure. Highly similar and conserved patterns were observed for predicted amino acid occupations in respective functional sites between the training and generated set (fig. 3C). Together with Shannon entropy profiles, these results demonstrated that PRO-LDM was able to identify and extract key evolutional information embedded in proteins from training sets and use them to design new sequences that resembled the native variants with key positions and residues retained, both in the aspect of protein scaffold and functions.

Both the amino acid composition and conformational states contribute to the functionality of proteins. Key residues accounting for the same function might be spatially adjacent in the folded protein, but being far apart in its primary sequence. Since our model had linear sequence inputs, we wondered if such global relationship in remote residue pairs can be grasped by the algorithm. The pairwise frequency distributions were calculated for each amino acid pair in all positions across sequences within MSAs. The correlation of these frequency distributions was determined in both training and PRO-LDM generated sets. As shown in fig. 3D, the amino acid pairwise relationships in both datasets exhibited remarkable similarity. PRO-LDM also surpassed the performance of the VAE model with higher average correlations (table S5). Furthermore, we examined whether generated sequences retained key functional domains suggested by previous experimental reports. Ten sequences from the generated set of MDH were randomly selected and inspected if two key domains (‘Ldh_1_N’ and ‘Ldh_1_C’) existed as shown in Pfam, each of which contained more than 100 amino acids and were far apart from each other (*46*). Both domains appeared in 9 out of 10 sequences, and only ‘Ldh_1_C’ domain showed in the last sequence. The results suggested that PRO-LDM generated new sequences preserved long-distance amino-acid relationships and key functional domains presenting in native proteins (fig. S1).

The diversity of newly generated sequences represents the capability of a deep learning model to explore beyond the known sequence space, which is a pivotal concern in the protein design field. Across the three datasets tested above, the diversity of sequences generated by PRO-LDM significantly exceeded those generated by VAE as well as the natural sets at the same sequence identity level within the cluster (fig. 3E). For instance, in the case of MDH dataset, the diversity of PRO-LDM generated sequences exceeded that of the VAE model and natural data by up to two-fold at 85% sequence identity. On the other hand, a stable protein structure was essential to maintain its functional conformation and enable effective biological functions. The *in vivo* stability of a protein was determined by the sequence order of certain amino acids (*47*), which could be estimated by the instability index from biopython with primary sequences. A value below 40 indicated a relatively high stability for the protein (*47*). The stability of PRO-LDM generated sequences was found to be equivalent to the training set for MDH and Luciferase_MSA (fig. 3F) with instability index value universally lower than 40. Yet the generated sequences were less stable than natural set in luciferase_RAW which may be due to its higher sequence diversity and length variety. Lastly, we determined the distribution of amino acid types for generated and natural sequences (fig. S2), which showed excellent agreement in all three datasets. Herein, it was demonstrated that PRO-LDM could unconditionally design protein sequences with higher diversity than the training datasets, while retaining the overall stability, evolutionary and physicochemical characteristics that defined proteins’ native structures and functions (fig. 3C, fig. S2).

### 4. PRO-LDM designs new proteins with tailored functional properties

Being minuscule molecular machines in living organisms, proteins undertook indispensable sets of biological functions (*48*). Yet the use of them in therapeutic and biomedical applications were often limited by the properties and functional performance of natural species (*49*). The tuning of natural protein properties, such as stability, solubility and physicochemical characterization, has been an important aspect of the protein design field with enormous practical potentials (*49, 50*). Since PRO-LDM was capable of grasping the characteristics of functional residues embedded in natural sequences and generating new variants with increased diversity, we further explored whether using labeled datasets combined with conditional diffusion module for semi-supervised training could enable the model to design new protein variants with tailored properties and functions.

The designated property or functional performance of a protein was denoted by a fitness value assigned during the training process. Proteins with high fitness excelled in specific screening experiments, showcasing enhanced thermal stability, more specific ligand binding capabilities, or achieving elevated expression levels in specific host organisms, *etc.* (*51*) Designing proteins with superior properties could be achieved by generating sequences with higher fitness values. The process was monitored to ensure effective generation in the model by recording changes in the property value of sequences during the iterations. Across all nine labeled datasets tested, the sequence fitness consistently exhibited a progressive approach towards the target value and tended to converge throughout the denoising process (fig.4A). This observation aligned with the sampling principle of the diffusion model, which initiated with random Gaussian noises and gradually eliminated them until a distribution was generated (fig.1A, fig.1B).

**Fig. 4.**
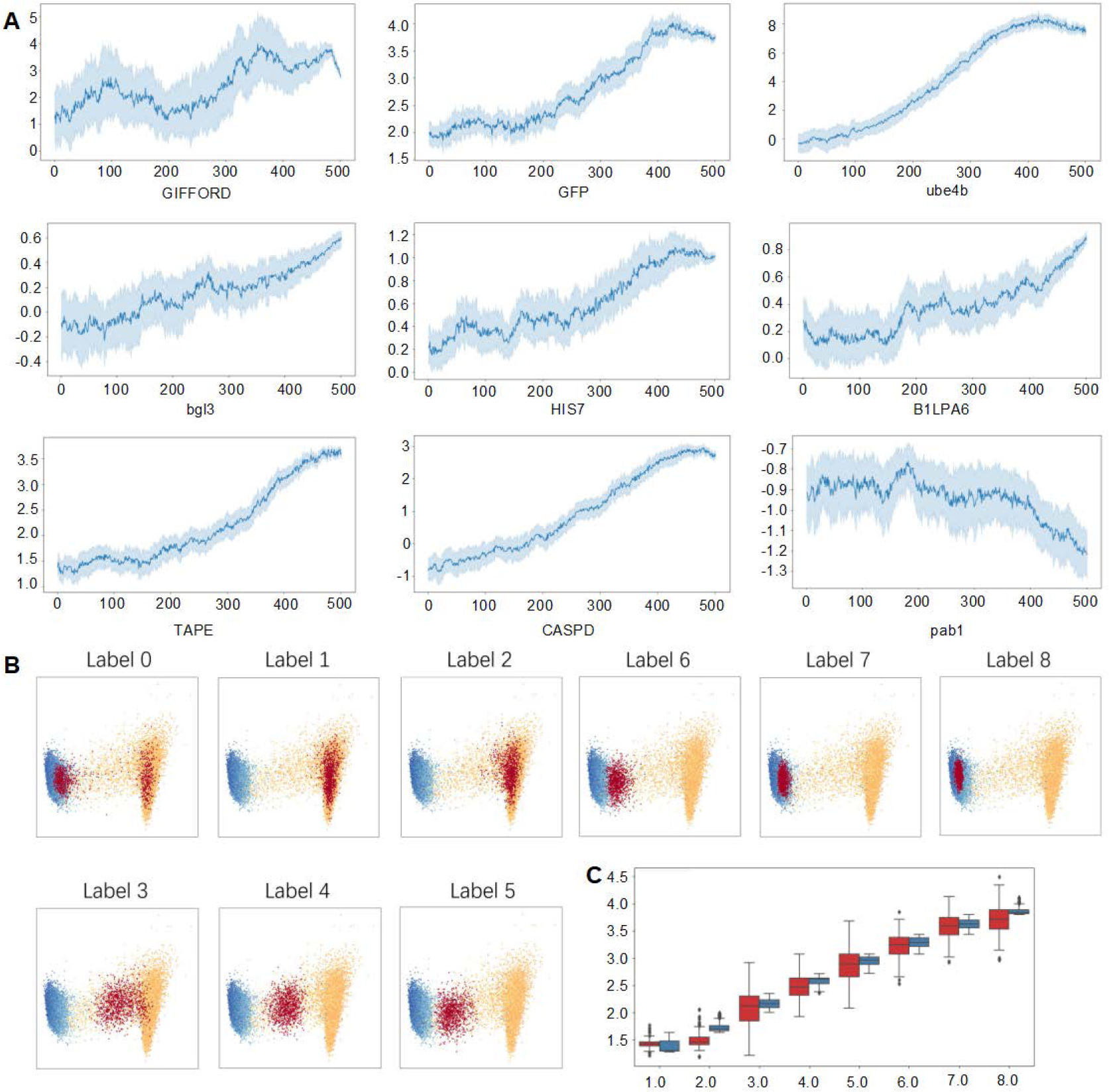
Conditional protein design with different labels. **(A)** The change of predicted fitness in conditional protein sequence generation. To visualize the convergence of protein fitness into the targeted area, high-fitness protein sequences were conditionally generated in 9 labeled datasets. The protein fitness was predicted using the latent variables generated during the denoising process, utilizing the pre-trained regressor. Sixty-four protein sequences were generated for each dataset. The intermediate dark blue line represents the average fitness value across 64 sequences. X axis: time step; Y axis: fitness. **(B)** Natural sequences of GFP dataset and conditionally generated protein sequences visualized in latent space. The sequences in the GFP dataset were divided into 8 labels based on their fitness, and sequences were conditionally generated for each label. The generated (red) and natural sequences (cool-color: higher fitness; warm-color: lower fitness) of each label were mapped into latent space and visualized using PCA. **(C)** Fitness distribution of natural and generated sequences for GFP dataset in each label. Sixty-four sequences of the GFP dataset were generated for each label, and their fitness was predicted using the regressor. The fitness distribution of the generated sequences (red) was compared to that of natural sequences (blue). X axis: label; Y axis: fitness.

In our model, protein variants with different levels of a specific property could be obtained by alteration in input labels. When the label was set to 0, new sequences were unconditionally generated with a distribution resembling that of the entire training set based on the visualization of their latent vectors, similar to using unlabeled datasets as described in the previous section. On the other hand, when a target value was assigned to the label, generated sequences exhibited clear alignment in their latent vector distribution against those with the same label in the training set. (fig.4B; fig. S3-10). Simultaneously, the regressor was employed to predict fitness value of generated protein sequences, which varied along with the input labels and reached levels comparable to corresponding sequences in the training data (fig.4C; fig. S11). PRO-LDM demonstrated capability of precisely adjusting specific protein properties by modifying the input label values.

Beyond the scope of generating new species in-distribution (ID) with those from the training sets, we further explore the potential of designing drastically different protein variants by sampling out of distribution (OOD) datapoints in the latent space, where generated data would have a relatively large discrepancy from the ID data. This method was used in OOD image generation and improved the generalization performance of ID tasks (*52*). Novel small molecules with enhanced properties in multiple domains could also be generated using an OOD-controlled diffusion model (*53*). Therefore, we referred to the classifier-free diffusion guidance in image generation and OOD sampling method to explore the relation and boundary between diversity and fitness distributions of samples in a directed-generation task.

In the classifier-free diffusion guidance (*54*), the sampling process is a linear combination of conditional and unconditional scores, as shown by Eq. (4). A hyperparameter *w* presented the strength of the classifier guidance, which consequently controlled variety and fidelity of generated samples (*54*). We herein evaluated different values of *w* between 0.1-1000. To compare our algorithm with SOTA models, we cross-referenced fitness prediction results on generated sequences from PRO-LDM regressor with Proteinbert (a self-supervised based deep language model for protein sequences), and Tranception (a transformer-based fitness prediction model leveraging autoregressive predictions and retrieval of homologous sequences at inference).

Similar to the sampling process in the classifier-free diffusion guidance in image generations, decreasing the strength of classifier guidance in PRO-LDM within a certain range enhanced the diversity of generated data but tended to decrease fidelity (*54*). When *w* was set between 0.1 to 1.0, we obtained all properly folded structures from the AlphaFold2 prediction (fig. 5A; fig. S12-13A). Decreasing *w* resulted in gradually increased diversity of generated protein sequences (table S5; fig. 5C; fig. S12-13C), accompanied by convergence of predicted fitness towards the vicinity of the targeted value (fig.5D-G; fig. S12-13). Using the GFP dataset as the example, designs precisely targeting a certain fitness could be achieved by setting *w* within a range less than 1, while decreasing the value of *w* enhanced the diversity of generated sequences. In contrast, when *w* was set between 1.0 and 10.0, we observed an incremental enhancement in the diversity of generated sequences (table S5), which still remained within the distribution of training data in the latent space (fig.5D).

**Fig. 5.**
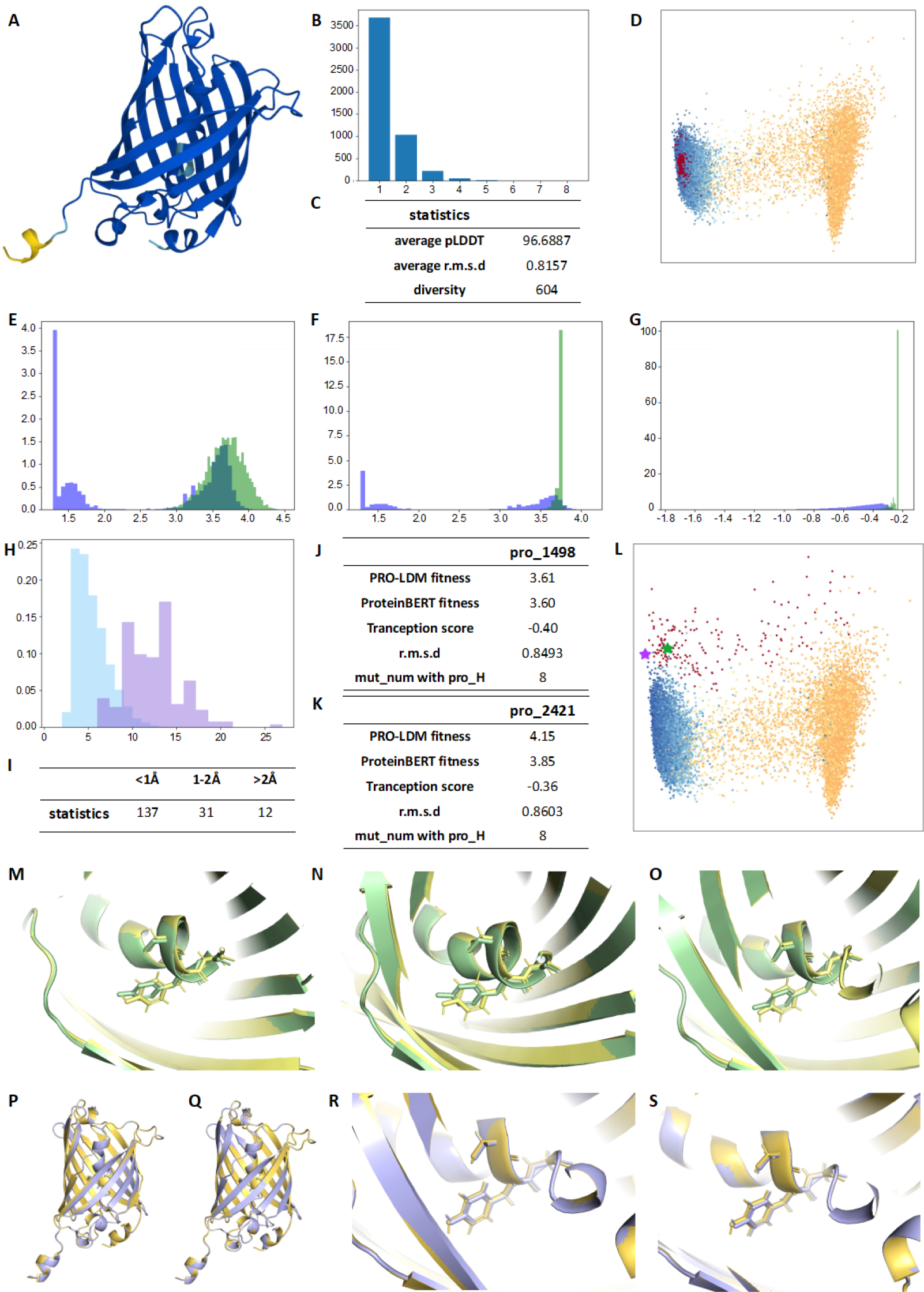
Analysis on OOD sequences with hyperparameter *w* = 1 and *w* = 20 for GFP dataset. (**A**) One of predicted structures for designed GFP protein. The color presented the pLDDT value of each site of amino acid. (**B**) The number of mutation sites of 5000 generated sequences. X axis: number of mutation sites; Y axis: the frequency of sequences. (**C**) The statistical data of the generated sequences. 5000 sequences were generated with hyperparameter set to 1. One hundred sequences were randomly selected and used to calculate the average pLDDT and r.m.s.d compared with protein with highest fitness in the training set. (**D**) Visualization of the latent space. The generated (red) and natural sequences (cool-color: higher fitness; warm-color: lower fitness) of each label were mapped into latent space and visualized by PCA. (**E-G**) Histogram of predicted fitness of generated sequences against fitness values in the training set. The value used in these three plots was predicted by three different platforms (E: PRO-LDM; F: ProteinBERT; G: Tranception). The x-axis of **E** and **G** represented sequence fitness values. X-axis of **G** represents the fitness score predicted by Tranception for all sequences. The Y-axis represented the relative number of sequences with the same fitness value (purple: natural sequences; green: generated sequences). (**H**) Histogram of mutation site counts for GFP training set and generated OOD samples when w setting to 20. X axis: number of mutation sites; Y axis: relative number of sequences. (**I**) The statistic of *r.m.s.d* value between the generated OOD samples and protein with highest fitness in the DMS dataset as the reference sequence. Out of 186 sequences, 137 had RMSD less than 1Å compared to pro_H, while 37 sequences had RMSD between 1 - 2Å, and 12 sequences had RMSD>2Å. (**J-K**) The predicted fitness/score, RMSD and mutation numbers of pro_1498 and pro_2421 against pro_H. (**L**) Visualization of the latent space for training set and the generated OOD sequences at w=20 (the green and purple stars represented pro_1498 and pro_2421, respectively). (**M-S**) Superimposition of predicted structures by Alphafold2. **M**, **N, O:** local visualization of three key residues composing chromophore between wt-GFP and three high fitness sequences (M: pro_H; N: pro_1498; pro_2421). **P**, **Q:** Superimposition of pro_H with pro_1498 (P) and pro_2421 (Q). **R**, **S**: local visualization of three key residues composing chromophore between pro_H and two generated proteins (R: pro_1498; S: pro_2421).

With *w* further increased to the range of 10-20, datapoints representing generated sequences gradually resided beyond the distribution of the training set (fig. S14D, S15D), leading to a significant increase in sequence diversity (table S5; fig. S14C, S15C), albeit with a decline in fitness predicted by ProteinBERT and Tranception (fig. S14E-G, S15E-G). Alphafold2 was employed to predict the structures for 100 randomly selected sequences out of a pool of 1000 total samples. The average pLDDT for predicted structures was 96.09 at *w* = 20 (fig. S15C), while 87 of 90 proteins showed a root mean square deviation (RMSD) < 2Å against the sequence with the highest fitness (pro_H) in the training and test dataset.

Subsequently, we focused on analyzing high fitness datapoints predicted by ProteinBERT at *w*=20 (label 7 and 8 predicted by our regressor). OOD datapoints were selectively filtered according to their distribution against the training set within the latent space, resulting in a total of 180 sequences (fig. 5L, red dots). Fig. 5H showed the number of mutations in these OOD sequences which was higher than that in the training set, while 168 out of 180 sequences had RMSD < 2 Å compared to the structure of pro_H (fig. 5I). Sequences with a Tranception score greater than -0.40 (the first quartile Tranception score of label 7 and 8 data in the trainset) were subsequently selected: pro_1498 (green star in fig. 5L; fig. 5J) and pro_2421 (purple star in fig. 5L; fig. 5K). Both sequences possessed high function-related score as predicted by all three platforms, with *RMSD* values less than 1 Å and 8 mutations compared with pro_H.

The fluorescence of GFP is influenced by the chromophore and its surrounding microenvironment (*55*). Therefore, we aligned the predicted structures of pro_1498 and pro_2421 with those of wild type(wt)-GFP and pro_H, respectively. The relative positions of chromophore in each protein, as well as the interactions between the chromophore and surrounding amino acids, were inspected. As shown in fig. 5M-O, there was a deviation angle between the phenyl rings of residue Y65 in pro_H, pro_1498, and pro_2421 compared to that in wt-GFP. However, the chromophores in pro_1498 and pro_2421 were perfectly superimposable with that in pro_H (fig. 5P-S). Such alignments suggested potentially superior fluorescence from generated sequences in accordance with experimentally verified mutants as compared to wt-GFP. From fig. S16, such improvement may be attributed to the elimination of hydrogen bonding between the hydroxyl group of Y65 at the chromophore and adjacent structural component on the β-barrel (H147, T202) in pro_H, pro_1498, and pro_2421, which was present in wt-GFP.

When *w* was further increased beyond 20, the generated data migrated towards the low fitness region within the latent space, accompanied by a substantial decrease in predicted fitness on ProteinBERT and Tranception (fig. S17D-G). This diminished the purpose of generating proteins with higher fluorescence with the current dataset and was not further pursued, but may be relevant in other property-tuning tasks, such as designing lower-solubility proteins for self-assembly (*56*). Moreover, when *w* reached values above 500, the generated sequences gradually lost their foldability and stability, resulting in a significant decline in pLDDT scores (fig. S18).

When the *w* value was set above 10, the distribution of fitness predicted by PRO-LDM regressor diverged gradually from those predicted by ProteinBERT and Tranception, with the magnitude of differences becoming more pronounced as *w* increased (fig. S14-16E-G). These differences may be related to the training data used. While PRO-LDM was trained on DMS data with limited generalization that induced overconfident predictions for OOD datapoints, both ProteinBERT and Tranception were pretrained on general large-scale protein databases such as UniRef. Yet such discrepancy may be alleviated by integrating PRO-LDM with other pretrained models thanks to its modular design. Our results also demonstrated that at a sweet spot of *w* = 20, PRO-LDM can generate OOD proteins with greater sequence diversity than mutation scanning experiments, without negating much of the structural and functional characteristics from the natural proteins. The designed variant may be subjected to further fitness iterative optimization by algorithms like ReLSO, to achieve superior functional performance from a novel base sequence.

In summary, PRO-LDM has demonstrated capabilities in directed protein design on its property and functions by leveraging sequence diversity against fitness fidelity through adjusting the guidance strength using hyperparameter *w*. This may enable the fine-tuning of protein species for designated performance under different scenarios.

## Discussion

The vast sequence space poses notable challenges in accurate protein design due to the astronomical possibility in combination of amino acids. Yet it also offers enormous opportunities particularly as the sequence length increases, leading to exponential growth in potentially foldable and functional new proteins. Deep learning algorithms offer an opportunity to explore beyond the human knowledgebase on these molecular machines without the complexity of traversing the whole sequence space, by learning intrinsic relationship embedded within amino acids and protein sequences from known species. These approaches are undisputedly gaining momentum as a prominent avenue for protein design, holding great promise for creating a broad spectrum of functional variants applicable in biomedicine and biotechnology.

There are three main categories of deep learning-based methodologies for protein sequence design. The first type, represented by unsupervised learning models like ProteinGAN (*45*), utilizes information from a specific protein family for training and generates novel functional sequences within that family. However, these models may not simultaneously optimize physicochemical properties, such as thermal stability, from a broader spectrum for the generated sequences. The second type, exemplified by directed optimization approaches like ReLSO (*38*), focuses on optimizing specific protein properties using labeled datasets of mutation data with corresponding fitness values, which are inherently limited by their reliance on labeled datasets. The third type adopts dedicated language models to train on genomic scale unlabeled protein sequences, which require substantial amount of data for training and incur high computational costs (*57, 58*).

In this work, we present a diffusion-based PRO-LDM model, aiming to identify commonalities and differences amongst amino acids in protein sequences, distinguish proteins with different functional performance within the latent space, and generate fine-tuned new sequences. The model utilizes a latent diffusion module founded on a jointly trained autoencoder, to effectively generates new sequences akin to natural protein, leveraging unsupervised learning for enhanced diversity and the ability to produce tailored functional variants under specific conditions. PRO-LDM excels in representation learning when compared to relatively primary ReLSO backbone. It exhibits proficiency in learning fundamental and functional representations from natural sequences, through comparisons of embeddings for amino acids with similar properties and visualizations of sequence representations within fitness landscapes. When employed in unconditional design, PRO-LDM generates sequences remarkably similar to natural proteins in terms of sequence identity, stability, local and global amino-acid relationships, but with enhanced diversity. This is attributed to diffusion models’ capability in adding noise to the input and training to reconstruct the original input without noise, which improves the resilience and versatility of the encoder in the model, to produce high-fidelity data resembling the native templates. In conditional protein design, PRO-LDM generates sequences with specific fitness labels, with clustering similar to corresponding natural sequences in the fitness landscape. Compared to fitness optimization algorithms in latent space such as ReLSO, PRO-LDM does not require complex norm-based negative sampling to achieve a convex latent space for gradient-based optimization. The process is also tunable using a hyperparameter *w*, where diversity against fitness fidelity can be tailored with respect to natural proteins that enables precision design in different applications. The generation of OOD protein sequences can be implemented by further increasing the value of *w* . These variants notably deviated from the distribution in the training dataset may substantially expand the exploration of sequence space and potentially be adopted to design new proteins with greater sequence diversity and distinguished functionality compared to the wild-types.

Beyond the current stage of work, PRO-LDM can benefit from training with more general and varieties of datasets. The deep mutation scanning data used in the current work underperformed in sequence diversity and distribution that limited model’s conditional generation capability. However, the introduction of more diverse dataset would require to process protein sequences with variable-lengths, which is challenging in the current setup by padding sequences to a fixed length. Model architecture design to accommodate these varieties in sequence data would enable more comprehensive training and further broaden the applicability of PRO-LDM. The applicability and effectiveness of the model can be further enhanced by but not limited to 1) latent space optimization techniques to explore and leverage the underlying latent representations; 2) combining with large-scale protein language models and multimodal learning to enable latent zero-shot/few-shot prediction and protein sequence generation.

Moreover, since PRO-LDM uses a modular design, its high integrability enables the model to utilize the pretraining or decoding modules in other SOTA deep-learning algorithms and further enhance the designing capability. For instance, in the current setup, we adopt the encoder module from ReLSO, which carries out effective conditional optimizations besides the capability of unsupervised protein design. The module can be interchanged with ESM-2 (*59*), which can allow the model to better comprehend the correlation between amino acid pairs and the structural details ingrained in the protein sequence. Enhanced precision in simulating the fitness landscape and better understanding of the structure-activity relationship allows more accurate design of novel species with desired property or functions that can lead to a wider range of applications. On the other hand, since PRO-LDM is a sequence-based protein design tool, combining it with structure-based generative models like RF-Diffusion can potentially provide an end-to-end pipeline for highly precise protein scaffold customization, by building a parallel neural network that cross-reference information from both aspects.

## Methods

### Neural network architecture

The backbone of our model was a jointly trained autoencoder as developed in ReLSO. The encoder employed a four-head transformer with six hidden layers, each having a dimension of 200, to capture the interactions between residues in the sequence. A bottleneck module, consisting of a fully connected layer, was applied to compress the embedded information into the latent space. The decoder used four 1D convolutional layers to reconstruct sequences from latent variables. Rectified linear unit’s (ReLU) activation and batch normalization layers were incorporated between convolutional layers, except for the final layer.

In parallel with the decoder, a MLP-based regressor, composed of one fully connected layer and featuring a dropout rate of 0.2, was employed to predict fitness in our model. We leveraged a classifier-free diffusion guidance model between the encoder and decoder, which consisted of a four-layer 1D-convolutional U-Net that captured the disturbed latent distribution of each diffusion step. This approach facilitated learning the latent space of the sequences and enabled the conditional generation of *z*.

### Network training and labeled fitness

The distinct labels were assigned to sequences based on various fitness ranges. An 8-label division method was adopted over rounding fitness values. The procedure entails preprocessing each dataset and visualizing the relationship between dataset length and fitness distribution. Boundaries for fitness and sequence length were then established and uniformly divided into eight segments, with a few datasets being divided into five segments. For the unsupervised task, the labels were uniformly set to 0.

### Network training and sequence generation

During the training stage, sequences were fed into the encoder as strings, using an embedding layer with a dimension of 100, followed by the transformer learning the interdependencies between residues. Subsequently, a bottleneck layer compressed the discrete, sparse, high-dimensional sequence information into a 64-dimensional latent variable *z*. All *z* values constituted a continuous, condensed latent space of sequences. The fitness label was simultaneously introduced into the conditional diffusion model as input to learn the distribution of information in the latent space. Finally, *z* served as input for both the decoder and regressor, where the fitness label was reintroduced during supervised learning, for sequence reconstruction and fitness prediction.

The model was trained for 500 epochs utilizing the AdamW optimizer with a cosine annealing learning rate starting at 0.00002 on four 32GB V100 graphics processing units and employing a batch size of 512 (*60*).

### Bioinformatic analysis of generated sequences

#### Dataset selection

##### Luciferase_RAW dataset

Alex *et al*. downloaded from sequences containing a luciferase-like domain (IPR011251) from InterPro database (https://www.ebi.ac.uk/interpro/) (*41*). The dataset comprised 69,130 sequences with a maximum sequence length of 504 amino acids. The sequence identity threshold used during the splitting of the training and validation sets was set at 70%. The dataset was used to evaluate model’s generation capability with training on homologous proteins within the same domain.

##### Luciferase_MSA dataset

Based on Luciferase_RAW dataset, Alex *et al*. used Clustal Omega with the profile Hidden Markov Model (HMM) of the bacterial luciferase family from Pfam to create a MSA version of Luciferase_RAW dataset (*41*). The aligned sequences were used to evaluate the impact of incorporating evolutionary information in the training dataset on the learning performance of PRO-LDM across various tasks.

##### MDH dataset

Donatas *et al*. constructed the MDH dataset utilizing a family of bacterial MDH (malate dehydrogenase) enzymes (*45*). The dataset had three characteristics: 1) it comprised 16,898 sequences, with an average length of 319 ± 18.2 amino acids; 2) the pairwise identity of the sequences was based on a threshold of 10%; 3) the identity threshold of sequence used during the splitting of the training and validation sets was set at 70%. MDH dataset was selected for dual validation of the unconditional generation capability of PRO-LDM and evaluate its generalization ability across diverse protein families.

##### Mutation datasets

We trained and tested conditional PRO-LDM on nine deep mutational scanning datasets: Gifford(*61*), GFP(*62*), TAPE(*63*), Bgl3(*64*), Pab1(*65*), Ube4b(*66*), HIS7(*67*), CAPSD(*68*), B1LPA6(*69*). The effect of substitutions was evaluated using the initial six datasets, while the remaining three datasets were employed to quantify the impact of insertions and deletions (indels). More details regarding these 9 datasets could be found in Table S2.

### Protein structure prediction by Alphafold2

Alphafold2 was used to predict structures for sequences generated during the denoising process (*20*). The service is provided for free by Zhejiang Gene Computation Platform (https://cloud.aigene.org.cn/).

### Amino acids’ Pearson’s correlation and dimension reduction

For each sequence randomly selected from dataset, a 2D matrix was generated to represent the likelihood of different amino acids occurring at each residue position in the matrix [20, seq_length]. Pearson’s correlation was then calculated to determine the relationship between each amino acid pair. The mean value was calculated from the amino acid embeddings of all sequences, followed by dimension reduction via PCA.

### Latent space representations of protein sequence visualization

To evaluate the capacity of sequence-level representation generated by PRO-LDM in the latent space and to distinguish the functionality of proteins, the sub-family accession numbers were obtained for luciferase from InterPro. The latent representation for the nine largest sub-family proteins were encoded and generated. PCA was then employed for dimension reduction and visualization of this representation into 2D space.

### Identity analysis of generated and natural sequences

Sequences (number same as training set) were generated at 50-epoch intervals for three unlabeled datasets (MDH, luciferase_MSA, luciferase_RAW) throughout the training process. BLAST (http://www.ncbi.nlm.nih.gov/BLAST/) was employed to conduct sequence alignments on the generated sequences and calculated their identities in comparison to the natural set, represented in box plots.

### Multiple sequence alignments and Shannon entropy

One thousand training sequences and 1,000 generated sequences were randomly selected, combined and input into Clustal Omega or alignment (*70*). In cases where a mismatch occurred at a specific site, the non-matching position was replaced with a dash (“-”), referred as a gap. The training and generated sequences were separated, and columns exhibiting over 75% gap ratio were removed. The Shannon entropies of both sets were calculated separately using the following equation:

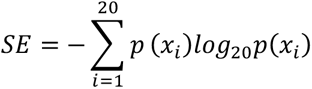

where *p*(*x*_*i*_ ) represented the frequency of amino acid i in one column of the MSA.

### Logo figure

One thousand sequences from natural and generated set were selected and input into Clustal Omega for alignment (*70*). Frequency matrices were computed for both sequence groups, where conserved positions were identified. TBtool and R were used for visualization. Both ggseqlogo (*71*) and ggplot2 (*72*) in R were employed to create and enhance the graphical presentations. The gridExtra package was used to consolidate the visualizations of both sequence groups and their correlation at the conserved positions.

### Pairwise amino acid frequency distribution

One thousand sequences were selected from both the training set and the generated set. Pairwise amino acid occurrence frequency matrices of dimensions [seq_length, seq_length] were computed for each sequence. The matrices were then reshaped to [1, seq_length × seq_length] and all sequences were concatenated in the training/generated set. Two groups of metrics were obtained with dimensions [seq_num, seq_length × seq_length], which were employed to calculate the Pearson’s correlation coefficient.

### Sequence diversity analysis

Generate sequences from PRO-LDM and VAE-based models in alignment with the number of training sets were used. MMseqs2 in MPI Bioinformatics Toolkit (https://toolkit.tuebingen.mpg.de/tools/mmseqs2) was used to cluster sequences at different identity threshold to get the diversity value and Origin was used to fit the curves using non-linear function (*73–75*).

### Sequence stability analysis

Sixty-four sequences were generated by PRO-LDM and VAE-based model respectively, and compared with randomly selected 64 sequences from the training sets. Biopython was used to calculate the instability index for each sequence, and matplotlib was used to make the box plot (*47*) .

### ProteinBERT and Tranception

ProteinBERT is a deep language model pretrained with Gene Ontology (GO) annotation prediction and tested with downstream tasks covering diverse protein properties (*76*). The pretrained ProteinBERT was fine-tuned using GFP dataset in TAPE (*77*) before testing PRO-LDM generated sequences. Tranception is a SOTA transformer-based fitness prediction model employed to test PRO-LDM generated sequences (*78*). A higher Tranception score indicated superior functionality.

### Average pLDDT and RMSD

When exploring the OOD samples under different *w* values, we randomly selected 100 sequences from the generated set and input them into Alphafold2 on Zhejiang Gene Computation Platform. We downloaded all successfully predicted PDB files, extracted the pLDDT value for each atom, and calculated the mean value for all atoms. Meanwhile, we used the Superimposer module in biopython to calculate the RMSD value between the generated protein and pro_H.

### Superimposition of predicted structures

The predicted structures for the generated protein, pro_H, and the wt-GFP by Alphafold2 were used. The structure pairs were aligned in PyMOL (https://pymol.org) to visualize the superimposition of chromophores with hydrogen bond visualized.

