## supporting information for "PRO-LDM: Protein Sequence Generation with a Conditional Latent Diffusion Model"

Sitao Zhang *et al.*

**This PDF file includes:**

Figs. S1 to S18
Tables S1 to S7
References (61, 66, 79 to 85)

**Table S1. Mann-Whitney U test of amino acids with similar and different biochemical properties.** Based on the amino acid types listed in the second column, the amino acid embeddings were separated into two distinct groups: a particular type of amino acids and all remaining amino acids. Following this, a Mann-Whitney U test was performed to determine the statistical significance of the observed differences between these two groups.

| | **Amino acid type** | **P-value** |
|---|---|---|
| Luciferase_MSA | hydrophobic aliphatic | 1.68e-08 |
| | hydrophobic aromatic | 2.68e-07 |
| | polar neutral | 1.13e-06 |
| | charge basic | 2.45e-05 |
| | charge acidic | 8e-04 |
| Luciferase_RAW | hydrophobic aliphatic | 2.58e-06 |
| | hydrophobic aromatic | 0.15 |
| | polar neutral | 1.66e-05 |
| | charge basic | 6.25e-05 |
| | charge acidic | 8e-04 |
| MDH | hydrophobic aliphatic | 1.04e-08 |
| | hydrophobic aromatic | 8.52e-07 |
| | polar neutral | 2.67e-06 |
| | charge basic | 5.20e-05 |
| | charge acidic | 8e-04 |

**Table S2. Description of Deep Mutational Scanning (DMS) datasets used in conditional PRO-LDM.**

| | Datasets | Description | Organism | Molecular function | length | Variants | Ref. |
|---|---|---|---|---|---|---|---|
| Substitutions | Gifford | CDR3 of human Immunoglobulin G antibodies | human | IgG binding | 20 | 90,459 | (*61*) |
| | GFP | Green fluorescent protein | *Aequorea victoria* | Fluorescence | 237 | 54,025 | (*79*) |
| | TAPE | Green fluorescent protein | *Aequorea victoria* | Fluorescence | 237 | 54,025 | (*80*) |
| | Bgl3 | β-glucosidase | *Streptococcus* sp. | Hydrolysis of β-glucosidic linkages | 501 | 26,653 | (*81*) |
| | Pab1 | RNA recognition motif (RRM) domain | *Saccharomyces cerevisiae* | Poly(A) binding | 75 | 40,852 | (*82*) |
| | Ube4b | Ubiquitination factor E4B U-box domain | *Mus musculus* | Ubiquitin-activating enzyme activity | 102 | 98,297 | (*66*) |
| Indels | HIS7 | Imidazoleglycerol-phosphate dehydratase | yeast | catalyzing the sixth step in the histidine biosynthesis pathway | 206-235 | 6,102 | (*83*) |
| | CAPSD | Capsid protein | Adeno-associated virus | delivery vectors for gene therapy | 735-749 | 250,907 | (*84*) |
| | B1LPA6 | Bifunctional chorismate mutase/prephenate dehydratase | - | Catalyzes the Claisen rearrangement and the decarboxylation/dehydration | 80-95 | 3,074 | (*85*) |

**Table S3. Fitness prediction metrics for DMS datasets.**

| Model | Dataset | Pearson'r | Spearman | m.s.e | L1 |
|---|---|---|---|---|---|
| PRO-LDM | Bgl3 | 0.49 | 0.44 | 1.52 | 0.97 |
| JT-AE | | 0.51 | 0.47 | 1.86 | 1.07 |
| PRO-LDM | GFP | 0.95 | 0.84 | 0.12 | 0.19 |
| JT-AE | | 0.90 | 0.81 | 0.33 | 0.37 |
| PRO-LDM | Pab1 | 0.43 | 0.39 | 3.19 | 1.51 |
| JT-AE | | 0.43 | 0.39 | 3.06 | 1.44 |
| PRO-LDM | Gifford | 0.82 | 0.47 | 0.26 | 0.38 |
| JT-AE | | 0.83 | 0.47 | 0.23 | 0.34 |
| PRO-LDM | TAPE | 0.87 | 0.65 | 0.95 | 0.88 |
| JT-AE | | 0.63 | 0.56 | 1.27 | 0.79 |
| PRO-LDM | Ube4b | 0.65 | 0.61 | 2.24 | 1.15 |
| JT-AE | | 0.60 | 0.56 | 2.89 | 1.30 |
| PRO-LDM | HIS7 | 0.83 | 0.69 | 0.07 | 0.17 |
| JT-AE | | 0.90 | 0.75 | 0.04 | 0.11 |
| PRO-LDM | CAPSD | 0.95 | 0.90 | 1.26 | 0.79 |
| JT-AE | | 0.94 | 0.88 | 1.33 | 0.80 |
| PRO-LDM | B1LPA6 | 0.58 | 0.51 | 0.12 | 0.27 |
| JT-AE | | 0.61 | 0.54 | 0.12 | 0.26 |

**Table S4. Correlation of Shannon entropy between natural and generated sequences.**

| Model | Dataset | Pearson'r | P-value | m.s.e |
|---|---|---|---|---|
| PRO-LDM | MDH | 0.9745 | 3.1653e-53 | 0.0253 |
| VAE | | 0.6689 | 1.1514e-64 | 0.2840 |
| PRO-LDM | Luciferase_MSA | 0.9921 | 8.6748e-20 | 0.0079 |
| VAE | | 0.9768 | 1.8404e-59 | 0.0226 |
| PRO-LDM | Luciferase_RAW | 0.7634 | 0.63 | 0.0851 |
| VAE | | 0.8832 | 1.1809e-53 | 0.0910 |

**Table S5: Average Pearson'r of amino-acid pair frequencies between natural and generated sequences.**

| Dataset | PRO-LDM | VAE |
|---|---|---|
| MDH | 0.9852 | 0.8504 |
| Luciferase_MSA | 0.8823 | 0.8296 |
| Luciferase_RAW | 0.8525 | 0.7970 |

**Table S6. Cluster numbers of 5000 generated sequences with different hyperparameter $w$ created by MMseq2.** For each hyperparameter w, 5000 sequences were generated and analyzed by MMseq2 using various similarity thresholds within clusters listed in the second to sixth columns of the first row. The table presented the quantity of clusters that indicated the diversity of the produced sequences.

| Sequence identity | 0 | 0.3 | 0.6 | 0.9 | 1 |
|---|---|---|---|---|---|
| $w = 0.1$ | 1 | 1 | 1 | 1 | 776 |
| $w = 0.2$ | 1 | 1 | 1 | 1 | 753 |
| $w = 0.3$ | 1 | 1 | 1 | 1 | 729 |
| $w = 0.4$ | 1 | 1 | 1 | 1 | 751 |
| $w = 0.5$ | 1 | 1 | 1 | 1 | 664 |
| $w = 0.6$ | 1 | 1 | 1 | 1 | 697 |
| $w = 0.7$ | 1 | 1 | 1 | 1 | 664 |
| $w = 0.8$ | 1 | 1 | 1 | 1 | 645 |
| $w = 0.9$ | 1 | 1 | 1 | 1 | 615 |
| $w = 1$ | 1 | 1 | 1 | 1 | 604 |
| $w = 2$ | 1 | 1 | 1 | 1 | 691 |
| $w = 3$ | 1 | 1 | 1 | 1 | 895 |
| $w = 4$ | 1 | 1 | 1 | 1 | 1330 |
| $w = 5$ | 1 | 1 | 1 | 1 | 1974 |
| $w = 10$ | 1 | 1 | 1 | 2 | 4694 |
| $w = 20$ | 1 | 1 | 1 | 1175 | 5000 |
| $w = 40$ | 1 | 1 | 10 | 4559 | 5000 |

**Table S7. Protein sequences for wild type GFP (P42212), pro_H (with highest fitness in DMS dataset), and two generated sequences when $w$ was set to 20. Mutation sites were highlighted.**

| Protein name | Protein sequence |
|---|---|
| wt-GFP (P42212) | SKGEELFTGVVPILVELDGDVNGHKFSVSGEGEGDATYGKLTLKFICTTGKLPVPWPTLVTTF<br>SYGVQCFSRYPDHMKQHDFFKSAMPEGYVQERTIFFKDDGNYKTRAEVKFEGDTLVNRIEL<br>KGIDFKEDGNILGHKLEYNYNSHNVYIMADKQKNGIKVNFKIRHNIEDGSVQLADHYQQN<br>TPIGDGPVLLPDNHYLSTQSALSKDPNEKRDHMVLLEFVTAAGITHGMDELYK |
| pro_H | SKGEELFTGVVPILVELDGDVNGHKFSVSGEGEGDASYGRLTLKFICTTGKLPVPWPTLVTTL<br>SYGVQCFSRYPDHMKQHDFFKSAMPEGYVQERTIFFKDDGSYKTRAEVKFEGDTLVNRIEL<br>KGIDFKEDGNILGHKLEYNYNSHNVYIMADKQKNGIKVNFKIRHNIEDGSVQLADHYQQN<br>TPIGDGPVLLPDNHYLSTQSALSKDPNEKRDHMVLLEFVTAAGITHGMDELYK |
| pro_1498 | SKGEELFTGVVPILVELDGDVNGHKFSVSGEGEGDATYGKLTLKFICTTGKLPVPWPTLVTTL<br>SYGVQCFSRYPDHMKQHDFFKSAMPEGYVQERTIFFKDDGNYRARAEVRFEGDTLVNRVE<br>LRGIDFKEDGNILGHKLEYNYNSHNVYIMADKQKNGIKVNFKIRHNIEDGSVQLADHYQQ<br>NTPIGDGPVLLPDNHYLSTQSALSKDPNEKRDHMVLLEFVTAAGITHGMDELYK |
| pro_2421 | SKGEELFTGVVPILVELDGDVNGHKFSVSGEGEGDATYGKLTLKFICTTGKLPVPWPTLVTTL<br>SYGVQCFSRYPDHMKQHDFFKSAMPEGYVQERTIFFKDDGNYKTRAEVKFEGDTLVNRIEL<br>KGIDFKEDGNILGHKLEYNFNSHNVYIMADKQKNGIKANFKIRHNIEDGSVQLADHYQRNT<br>PIGDGPVLLPDNHYLSTQSVLSKDPNEERDHMVLLEFVTAAGITHGMDELYK |

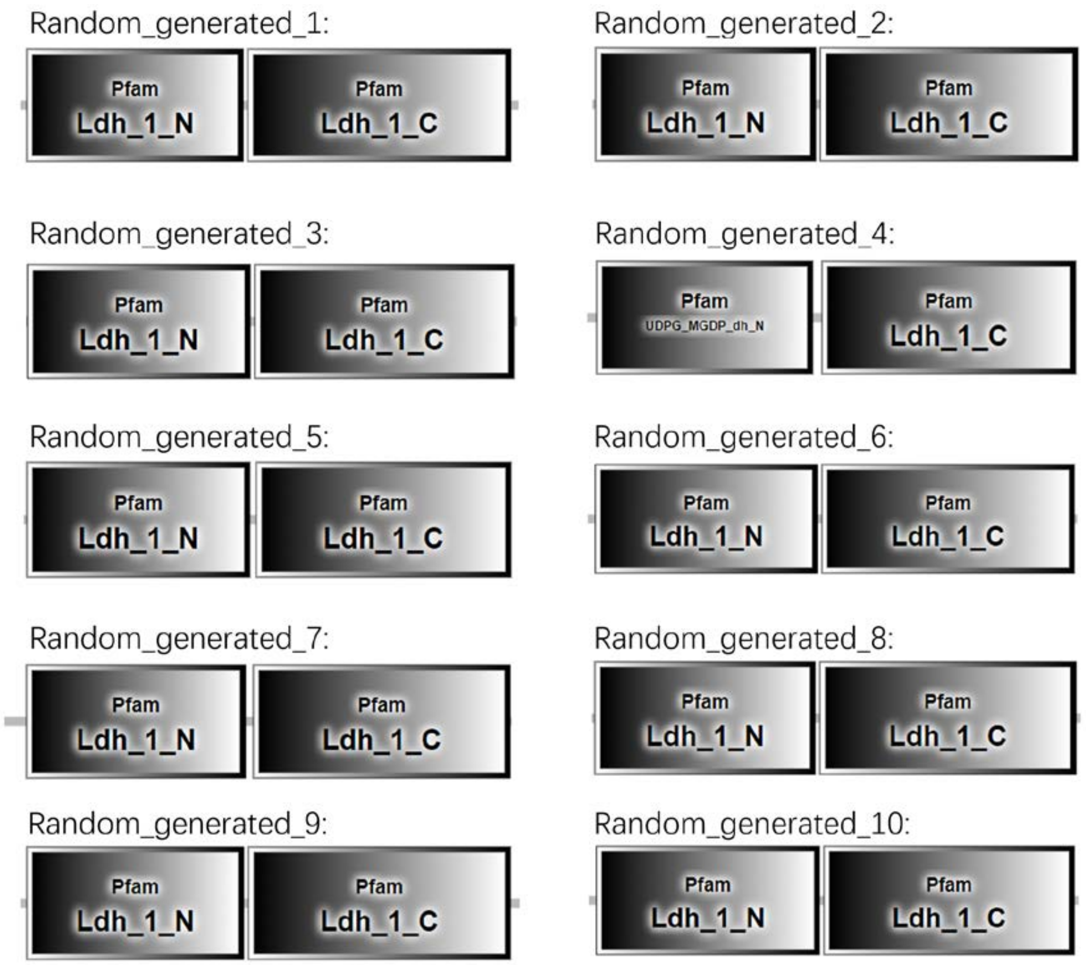


**Fig. S1. Pfam domains of 10 randomly generated sequences for MDH dataset.** Domain 'Ldh_1_N' and 'Ldh_1_C' contained more than 100 amino acids and were far apart from each other. 'Ldh_1_N' is the NAD binding domain and 'Ldh_1_C' is the α/β C-terminal domain of lactate/malate dehydrogenase. Nine out of ten generated sequences contained both of these two domains.

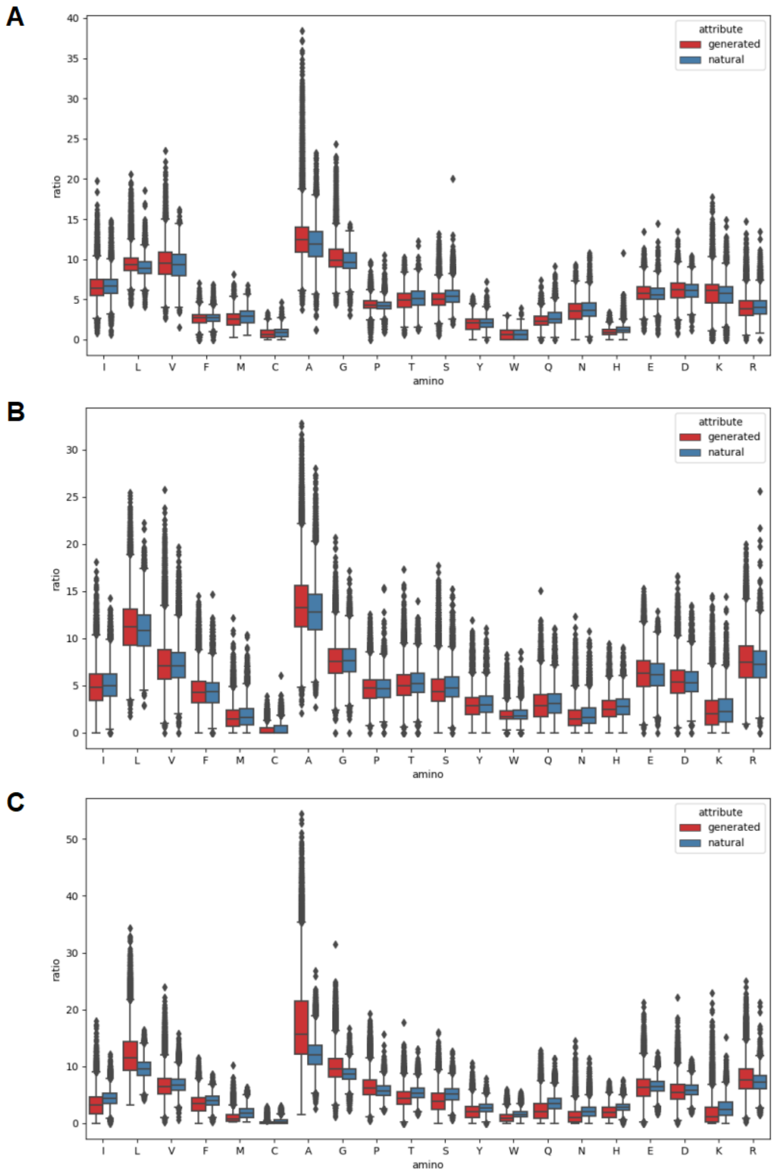


**Fig. S2. Amino acid composition of natural and generated protein sets**. **(A)** MDH; **(B)** Luciferase_MSA; **(C)** Luciferase_RAW.

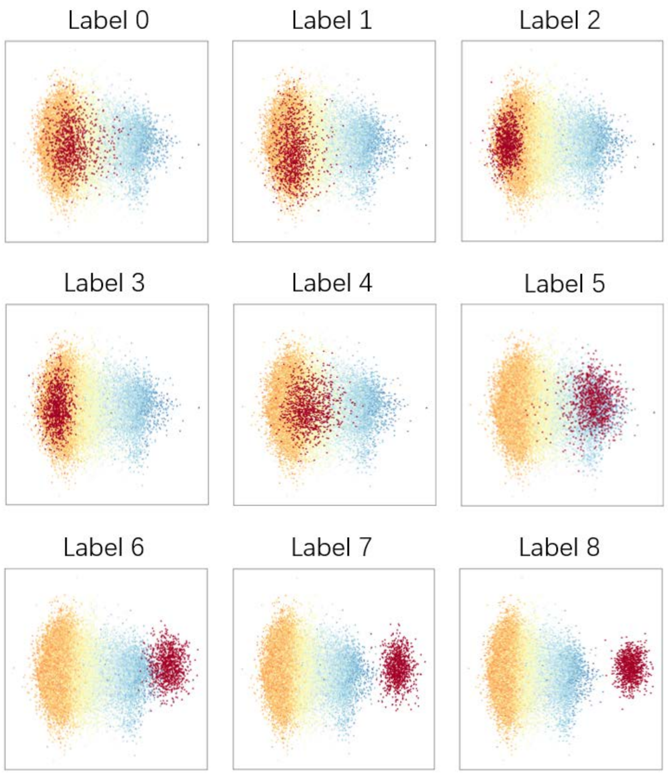


**Fig. S3. Natural sequences (cool-color: higher fitness; warm-color: lower fitness) and conditionally generated protein sequences (red) visualized in latent space (Gifford).**

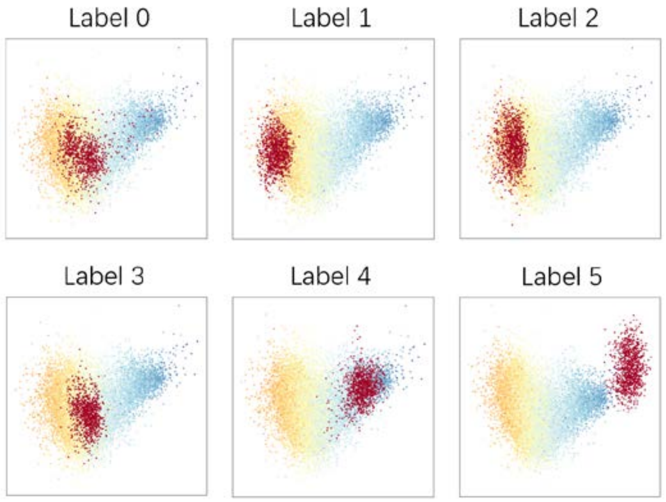


**Fig. S4. Natural sequences (cool-color: higher fitness; warm-color: lower fitness) and conditionally generated protein sequences (red) visualized in latent space (Ube4b).**

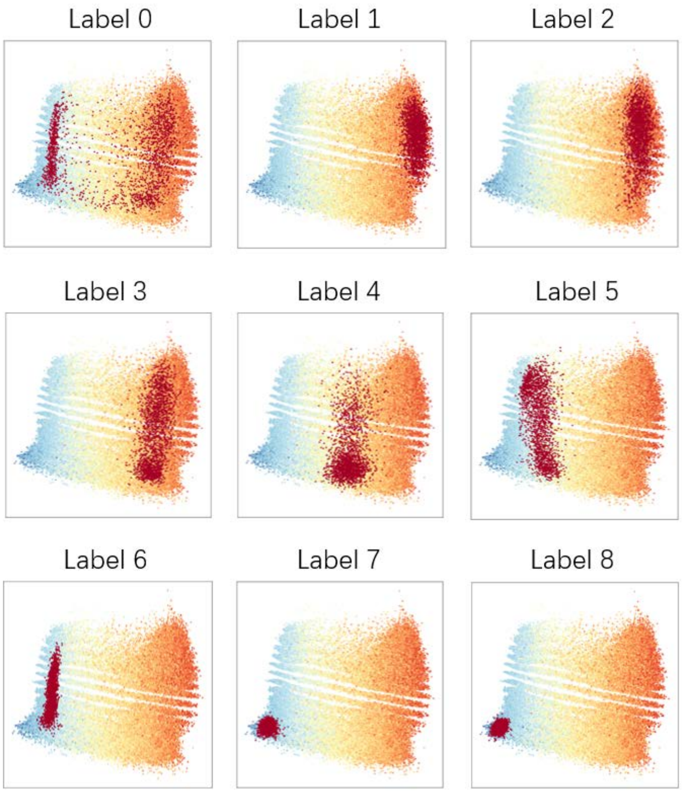


**Fig. S5. Natural sequences (cool-color: higher fitness; warm-color: lower fitness) and conditionally generated protein sequences (red) visualized in latent space (CAPSD).**

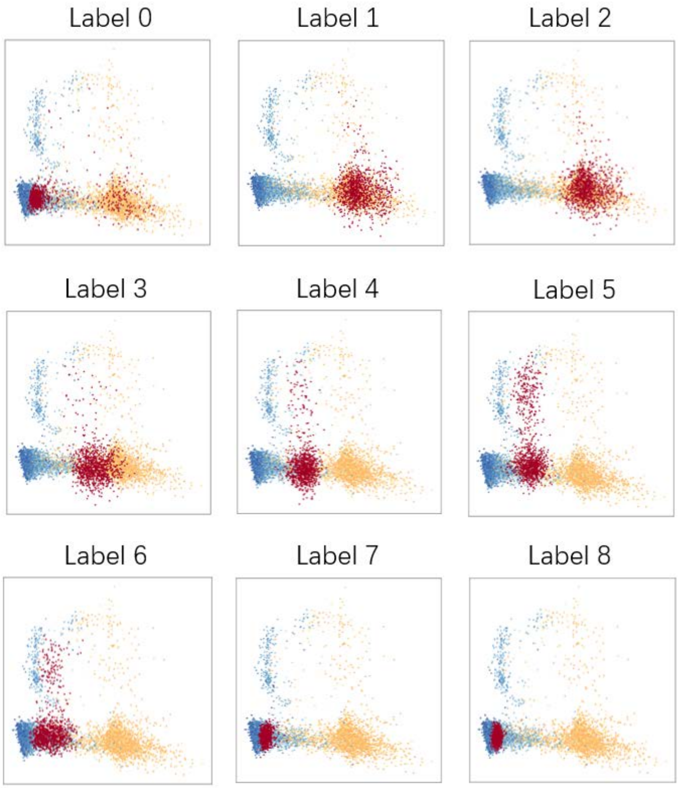


**Fig. S6. Natural sequences (cool-color: higher fitness; warm-color: lower fitness) and conditionally generated protein sequences (red) visualized in latent space (TAPE).**

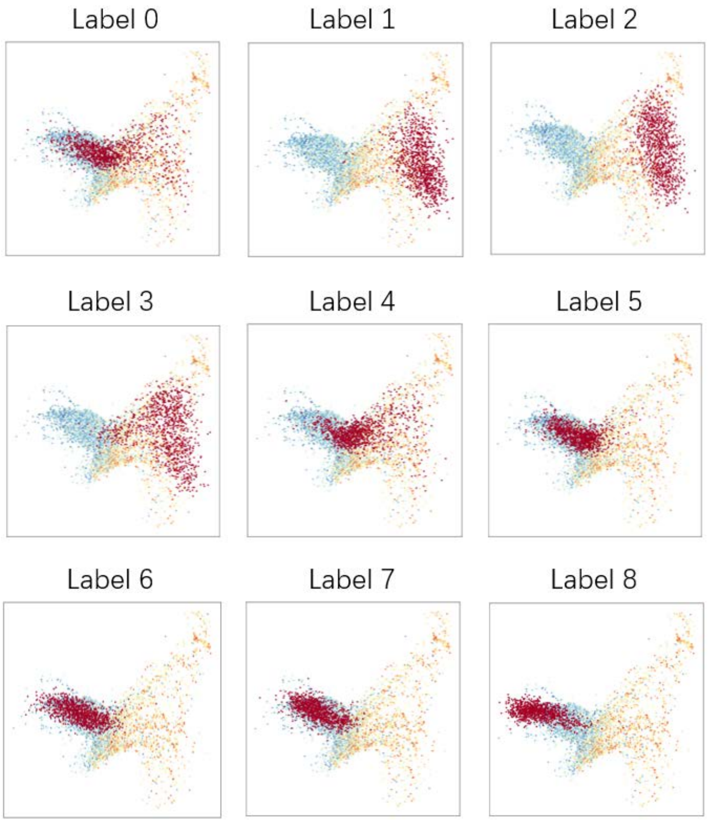


**Fig. S7. Natural sequences (cool-color: higher fitness; warm-color: lower fitness) and conditionally generated protein sequences (red) visualized in latent space (Bgl3).**

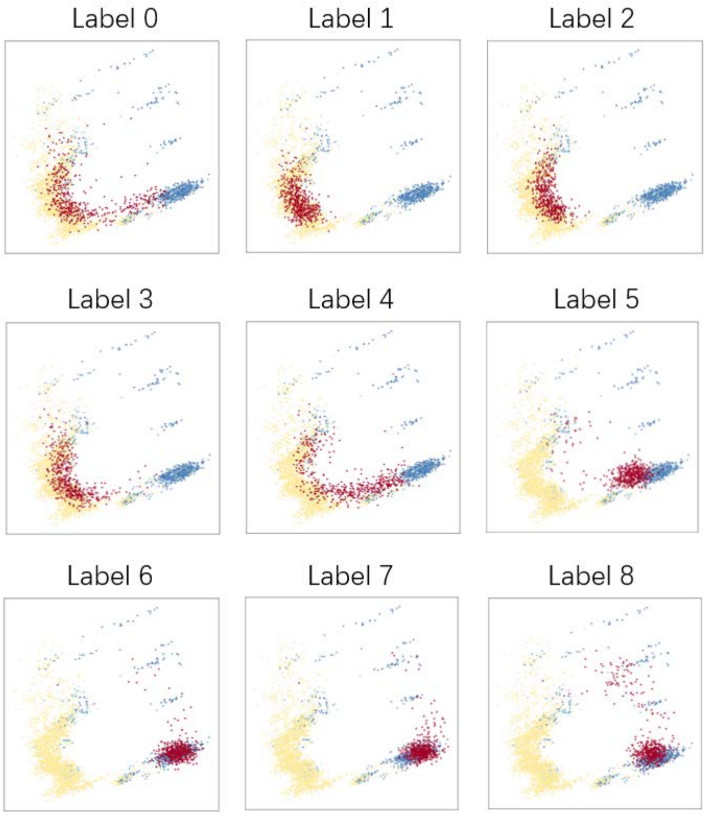


**Fig. S8. Natural sequences (cool-color: higher fitness; warm-color: lower fitness) and**

**conditionally generated protein sequences (red) visualized in latent space (HIS7).**

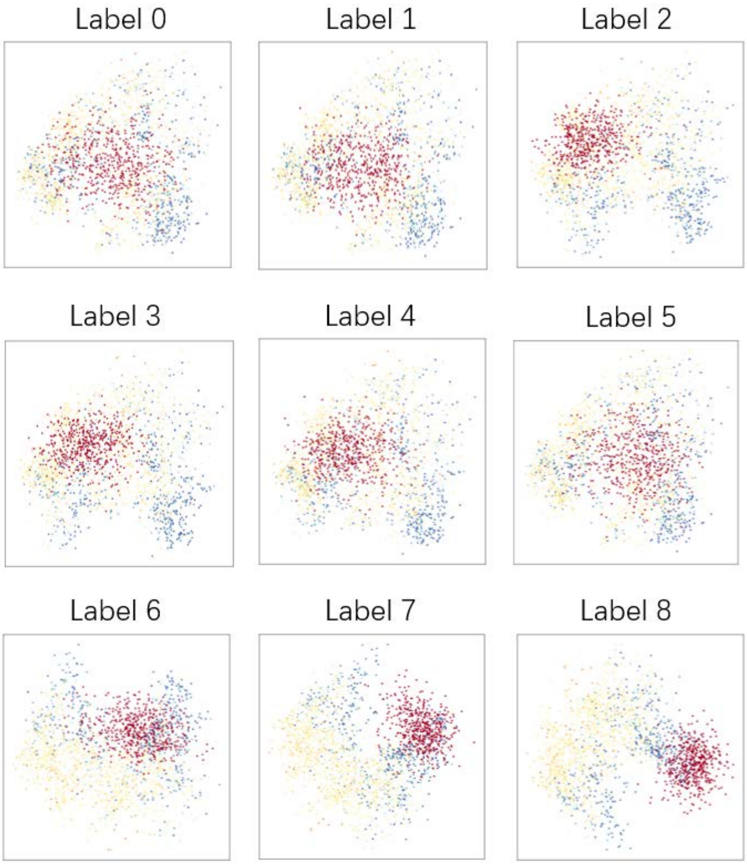


**Fig. S9. Natural sequences (cool-color: higher fitness; warm-color: lower fitness) and conditionally generated protein sequences (red) visualized in latent space (B1LPA6).**

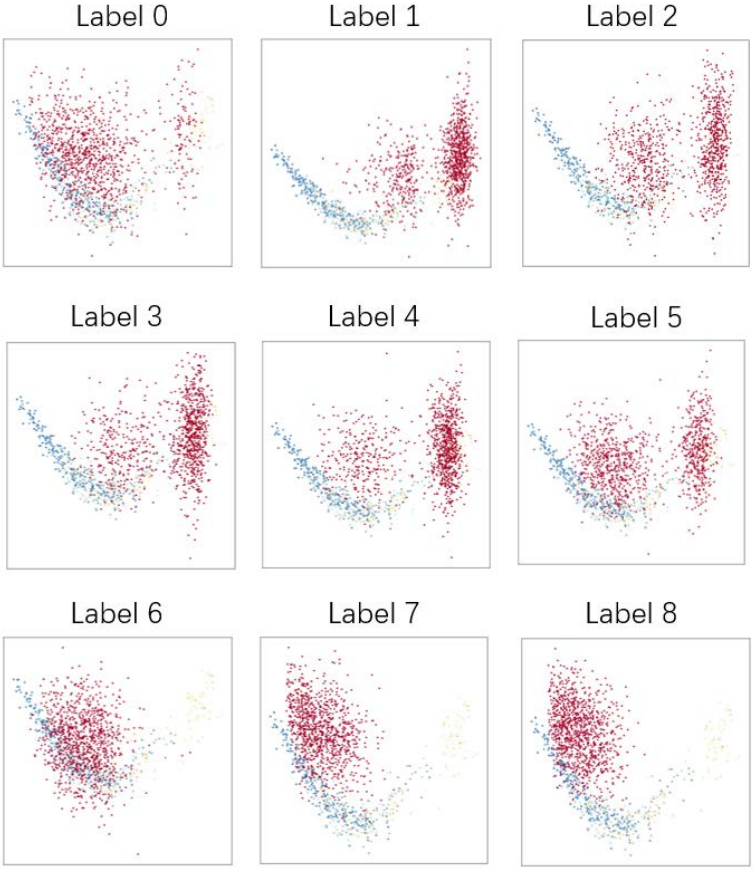


**Fig. S10. Natural sequences (cool-color: higher fitness; warm-color: lower fitness) and conditionally generated protein sequences (red) visualized in latent space (Pab1).**

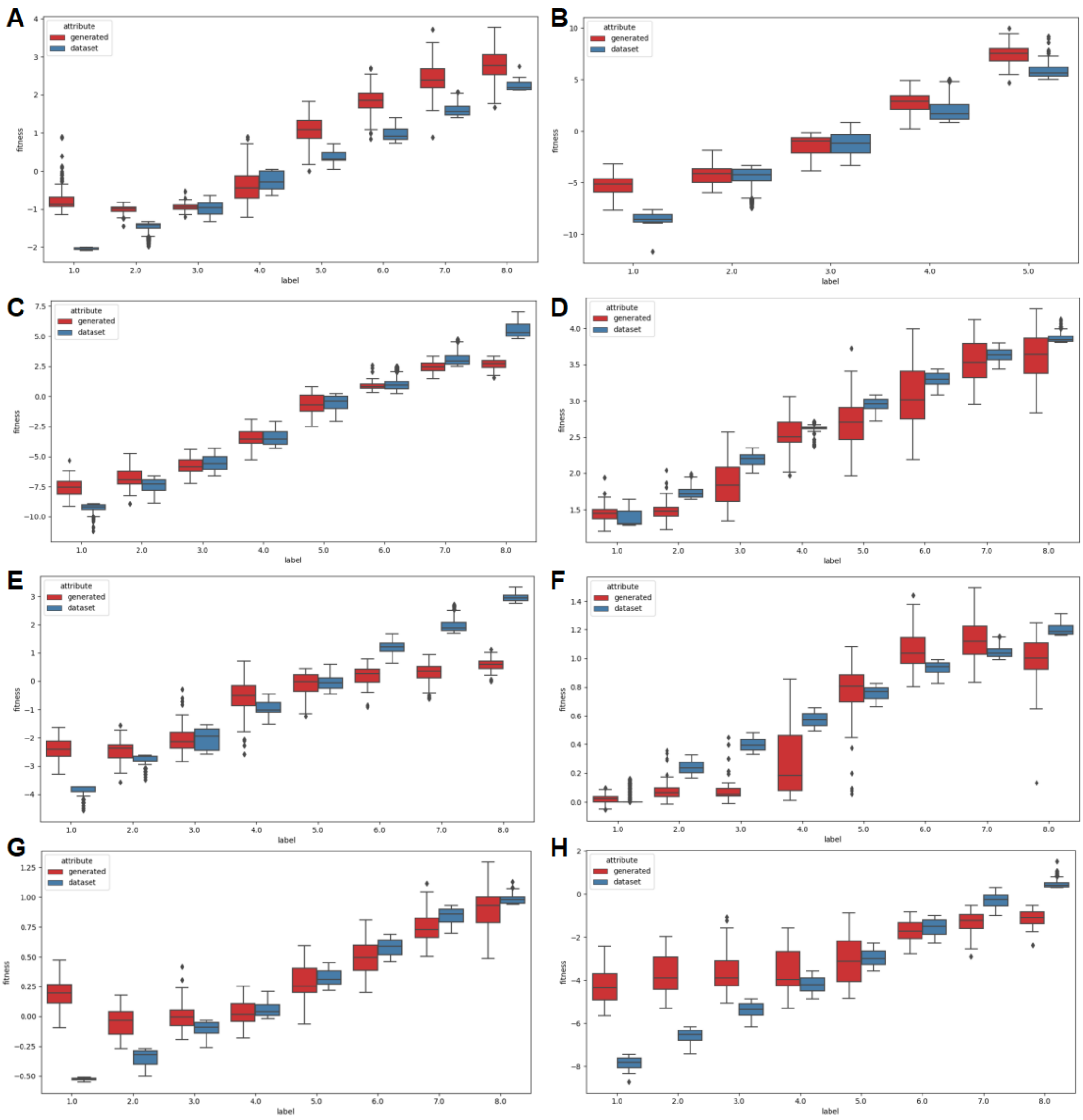


**Fig. S11. Fitness distribution of natural (blue) and generated (red) sequences of 8 mutation datasets in each label. (A)** Gifford; **(B)** Ube4b; **(C)** CAPSD; **(D)** TAPE; **(E)** Bgl3; **(F)** HIS7; **(G)** B1LPA6; **(H)** Pab1.

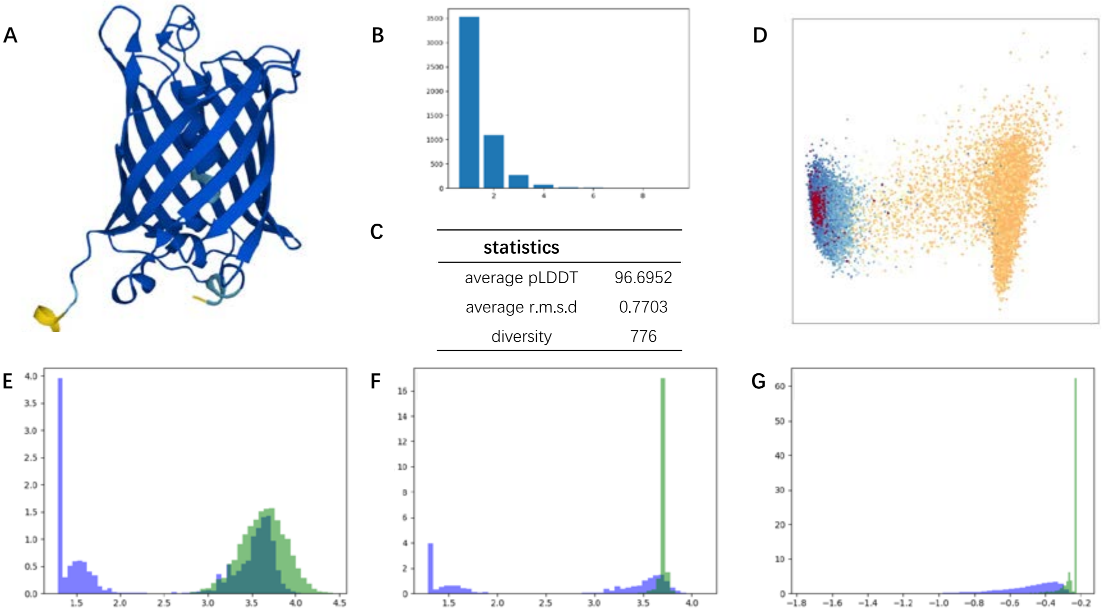


**Fig. S12. Analysis of generated sequences with hyperparameter $w$ = 0.1 for GFP dataset. (A)** An example of predicted structures for designed GFP protein. The color presented the pLDDT value of each site of amino acid (deep blue: pLDDT > 90; light blue: 90 > pLDDT > 70; yellow: 70 > pLDDT > 50). **(B)** The number of mutation sites for 5000 generated sequences. **(C)** The statistical data of the generated sequences. 5000 sequences were generated with hyperparameter $w$ set to 0.1. One hundred sequences were randomly selected and used to calculate the average pLDDT and RMSD **(D)** Visualization of the latent space. The generated (red) and natural sequences (cool-color: high fitness; warm-color: low fitness) of each label were mapped into the latent space and visualized using PCA. **(E-G)** Histogram of predicted fitness of generated sequences against fitness values in the training set. The value used in these plots was predicted by three different platforms (**E**, PRO-LDM; **F**, ProteinBERT; **G**, Tranception). The X-axis of **E** and **G** represented sequence fitness values. X-axis of **G** represents the fitness score predicted by Tranception for all sequences. The Y-axis of the subplots represented the relative number of sequences with the same fitness value (purple: natural sequences; green: generated sequences).

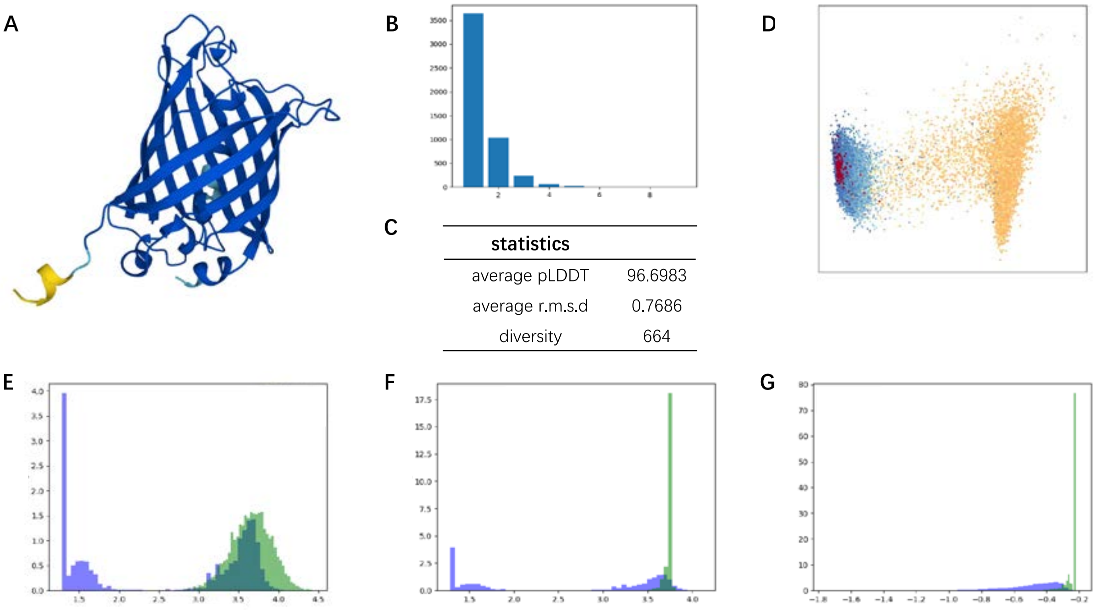


**Fig. S13. Analysis with hyperparameter $w$ = 0.5 for GFP dataset. (A)** An example of predicted structures for designed GFP protein. The color presented the pLDDT value of each site of amino acid (deep blue: pLDDT > 90; light blue: 90 > pLDDT > 70; yellow: 70 > pLDDT > 50). **(B)** The number of mutation sites for 5000 generated sequences. **(C)** The statistical data of the generated sequences. 5000 sequences were generated with hyperparameter $w$ set to 0.5. One hundred sequences were randomly selected and used to calculate the average pLDDT and RMSD **(D)** Visualization of the latent space. The generated (red) and natural sequences (cool-color: high fitness; warm-color: low fitness) of each label were mapped into the latent space and visualized using PCA. **(E-G)** Histogram of predicted fitness of generated sequences against fitness values in the training set. The value used in these plots was predicted by three different platforms (**E**, PRO-LDM; **F**, ProteinBERT; **G**, Tranception). The X-axis of **E** and **G** represented sequence fitness values. X-axis of **G** represents the fitness score predicted by Tranception for all sequences. The Y-axis of the subplots represented the relative number of sequences with the same fitness value (purple: natural sequences; green: generated sequences).

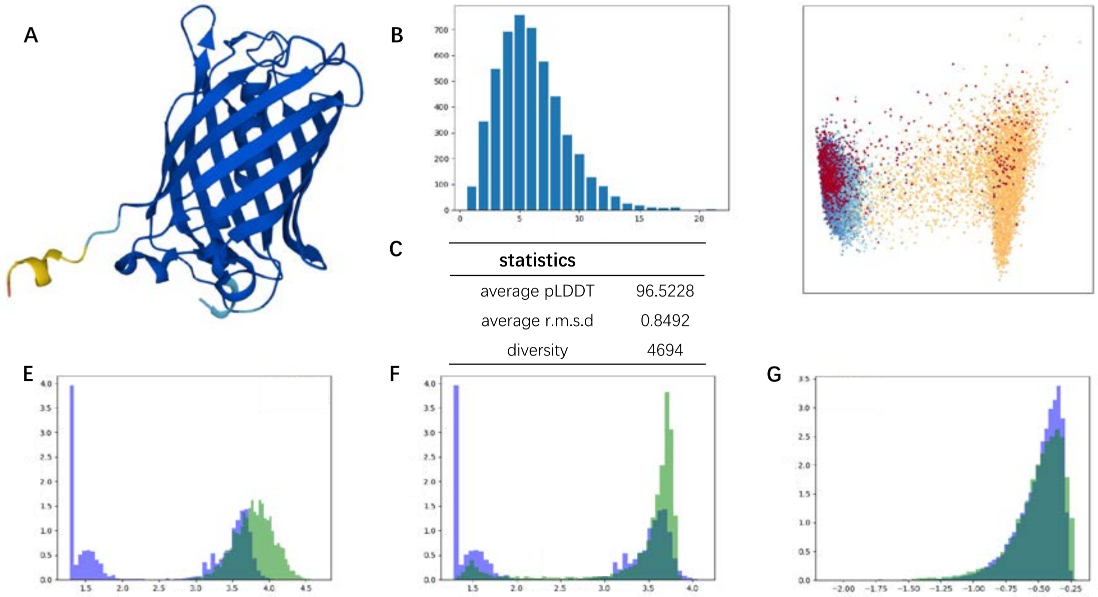


**Fig. S14. Analysis with hyperparameter $w$ = 10 for GFP dataset. (A)** An example of predicted structures for designed GFP protein. The color presented the pLDDT value of each site of amino acid (deep blue: pLDDT > 90; light blue: 90 > pLDDT > 70; yellow: 70 > pLDDT > 50). **(B)** The number of mutation sites for 5000 generated sequences. **(C)** The statistical data of the generated sequences. 5000 sequences were generated with hyperparameter $w$ set to 10. One hundred sequences were randomly selected and used to calculate the average pLDDT and RMSD **(D)** Visualization of the latent space. The generated (red) and natural sequences (cool-color: high fitness; warm-color: low fitness) of each label were mapped into the latent space and visualized using PCA. **(E-G)** Histogram of predicted fitness of generated sequences against fitness values in the training set. The value used in these plots was predicted by three different platforms (**E**, PRO-LDM; **F**, ProteinBERT; **G**, Tranception). The X-axis of **E** and **G** represented sequence fitness values. X-axis of **G** represents the fitness score predicted by Tranception for all sequences. The Y-axis of the subplots represented the relative number of sequences with the same fitness value (purple: natural sequences; green: generated sequences).

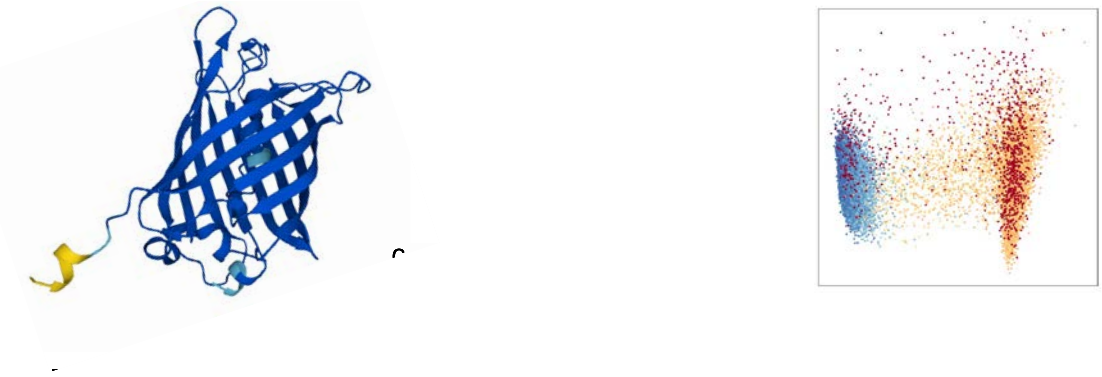

**Fig. S15. Analysis with hyperparameter $w$ = 20 for GFP dataset. (A)** An example of predicted structures for designed GFP protein. The color presented the pLDDT value of each site of amino acid (deep blue: pLDDT > 90; light blue: 90 > pLDDT > 70; yellow: 70 > pLDDT > 50). **(B)** The number of mutation sites for 5000 generated sequences. **(C)** The statistical data of the generated sequences. 5000 sequences were generated with hyperparameter $w$ set to 20. One hundred sequences were randomly selected and used to calculate the average pLDDT and RMSD **(D)** Visualization of the latent space. The generated (red) and natural sequences (cool-color: high fitness; warm-color: low fitness) of each label were mapped into the latent space and visualized using PCA. **(E-G)** Histogram of predicted fitness of generated sequences against fitness values in the training set. The value used in these plots was predicted by three different platforms (**E**, PRO-LDM; **F**, ProteinBERT; **G**, Tranception). The X-axis of **E** and **G** represented sequence fitness values. X-axis of **G** represents the fitness score predicted by Tranception for all sequences. The Y-axis of the subplots represented the relative number of sequences with the same fitness value (purple: natural sequences; green: generated sequences).

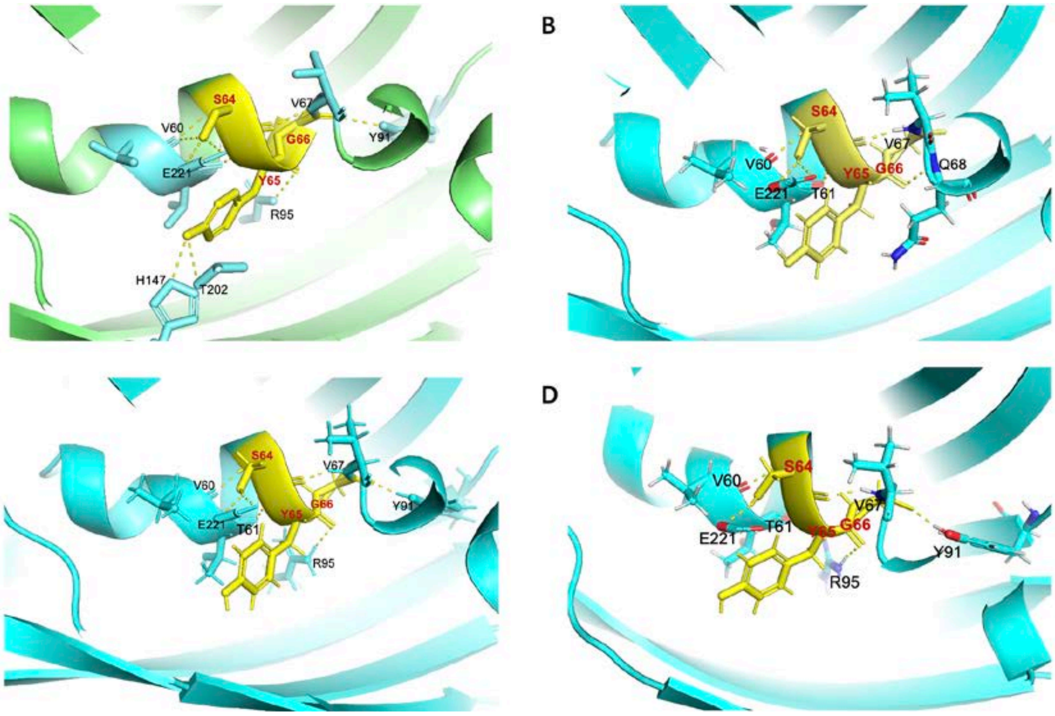


**Fig. S16. Visualization of hydrogen bond around chromophore. (A)** wt-GFP (P42212). **(B)** pro_H (the protein with highest fitness in the DMS dataset). **(C)** pro_1498. **(D)** pro_2421.

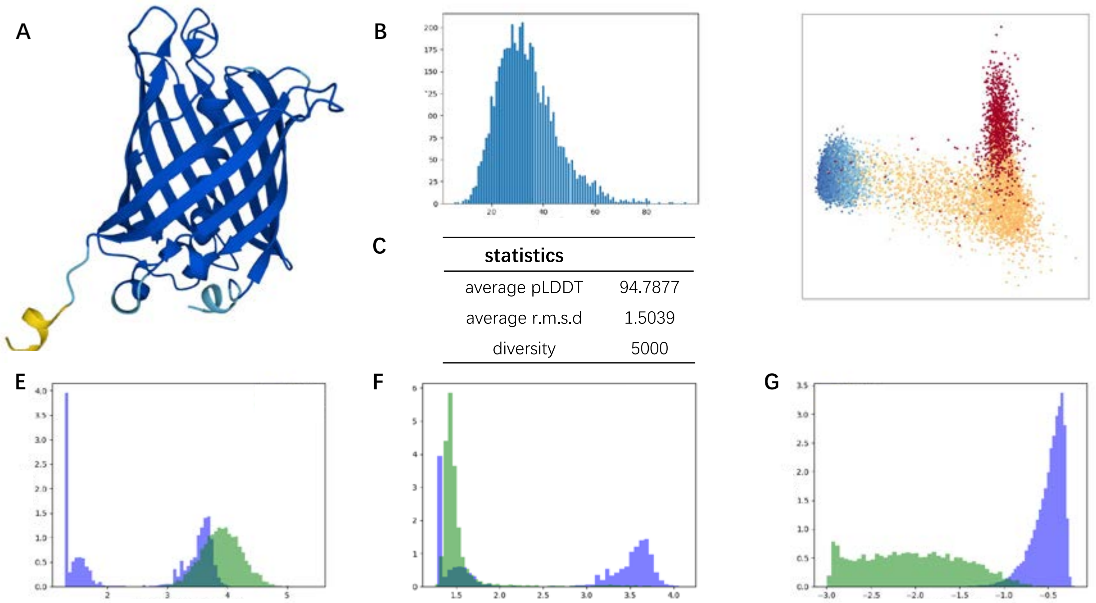


| statistics | |
|---|---|
| average pLDDT | 94.7877 |
| average r.m.s.d | 1.5039 |
| diversity | 5000 |

**Fig. S17. Analysis with hyperparameter $w$ = 40 for GFP dataset. (A)** An example of predicted structures for designed GFP protein. The color presented the pLDDT value of each site of amino acid (deep blue: pLDDT > 90; light blue: 90 > pLDDT > 70; yellow: 70 > pLDDT > 50). **(B)** The number of mutation sites for 5000 generated sequences. **(C)** The statistical data of the generated sequences. 5000 sequences were generated with hyperparameter $w$ set to 40. One hundred sequences were randomly selected and used to calculate the average pLDDT and RMSD **(D)** Visualization of the latent space. The generated (red) and natural sequences (cool-color: high fitness; warm-color: low fitness) of each label were mapped into the latent space and visualized using PCA. **(E-G)** Histogram of predicted fitness of generated sequences against fitness values in the training set. The value used in these plots was predicted by three different platforms (**E**, PRO-LDM; **F**, ProteinBERT; **G**, Tranception). The X-axis of **E** and **G** represented sequence fitness values. X-axis of **G** represents the fitness score predicted by Tranception for all sequences. The Y-axis of the subplots represented the relative number of sequences with the same fitness value (purple: natural sequences; green: generated sequences).

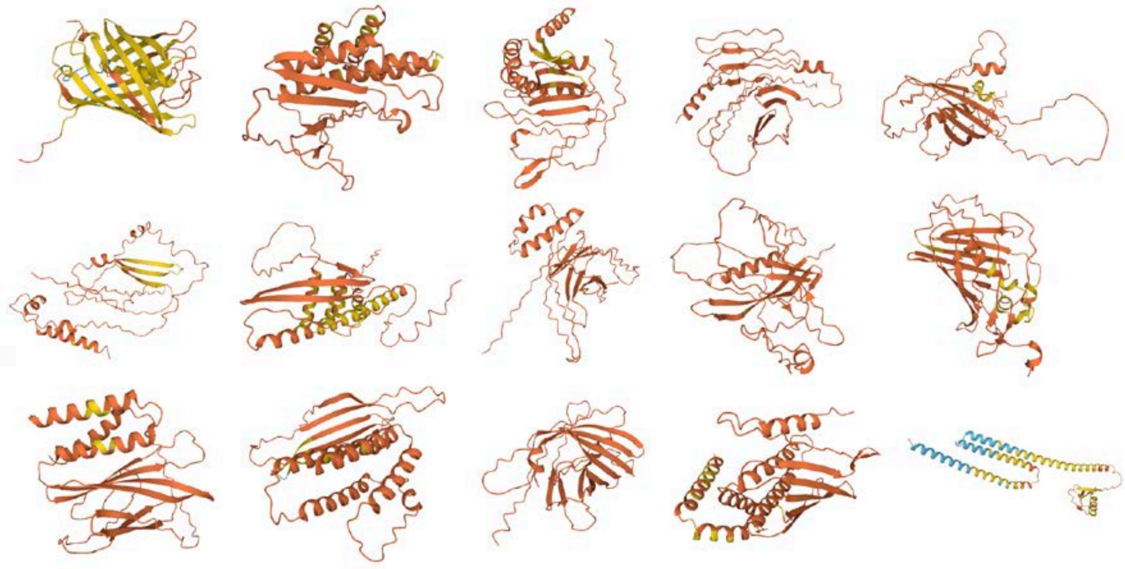

**Fig. S18. Predicted structures of generated sequences with hyperparameter $w$ = 1000 for GFP dataset.**
